# Neuraminidase 1 regulates the cellular state of microglia by modulating the sialylation of Trem2

**DOI:** 10.1101/2024.05.20.595036

**Authors:** Leigh Ellen Fremuth, Huimin Hu, Diantha van de Vlekkert, Ida Annunziata, Jason Andrew Weesner, Elida Gomero, Alessandra d’Azzo

**Affiliations:** Department of Genetics, St. Jude Children’s Research Hospital, Memphis, TN, 38105 USA; Compliance Office, St. Jude Children’s Research Hospital, Memphis, TN, 38105 USA; Department of Anatomy and Neurobiology, College of Graduate Health Sciences, University of Tennessee Health Science Center, Memphis, TN 38163, USA

**Keywords:** Neuraminidase 1, Trem2, Sialylation, Microglia, Neuroinflammation, Sialidosis, Alzheimer disease, Neurodegeneration

## Abstract

Neuraminidase 1 (Neu1) cleaves terminal sialic acids from sialoglycoproteins in endolysosomes and at the plasma membrane. As such, Neu1 regulates immune cells, primarily those of the monocytic lineage. Here we examined how Neu1 influences microglia by modulating the sialylation of full-length Trem2 (Trem2-FL), a multifunctional receptor that regulates microglial survival, phagocytosis, and cytokine production. When Neu1 was deficient/downregulated, Trem2-FL remained sialylated, accumulated intracellularly, and was excessively cleaved into a C-terminal fragment (Trem2-CTF) and an extracellular soluble domain (sTrem2), enhancing their signaling capacities. Sialylated Trem2-FL (Sia-Trem2-FL) did not hinder Trem2-FL–DAP12–Syk complex assembly but impaired signal transduction through Syk, ultimately abolishing Trem2-dependent phagocytosis. Concurrently, Trem2-CTF–DAP12 complexes dampened NFκB signaling, while sTrem2 propagated Akt-dependent cell survival and NFAT1-mediated production of TNFα and CCL3. Because Neu1 and Trem2 are implicated in neurodegenerative/neuroinflammatory diseases, including Alzheimer disease (AD) and sialidosis, modulating Neu1 activity represents a therapeutic approach to broadly regulate microglia-mediated neuroinflammation.

## Introduction

Glycosidic posttranslational modifications on membrane glycoproteins influence their ability to interact with other moieties, respond to cues, and modulate downstream signaling events.^1^ *N*-acetylneuraminic (sialic) acids play a pivotal role, given their negative charge, terminal position on glycan chains, and ubiquitous distribution.^2^ The extent of sialylation of glycoconjugates is controlled by sialyltransferases, which add sialic acids, and sialidases/neuraminidases 1-4 (Neu1-4), which remove them.^3^ Neu1 is the most ubiquitous of the mammalian sialidases, functioning in lysosomes and at the plasma membrane (PM).^4,5^ Within lysosomes, Neu1 initiates the catabolism of glycoconjugates by removing their terminal sialic acids.^6^ At the PM, it modulates the sialic acid content of glycoprotein receptors, influencing their functions.^6,7^ In humans, Neu1 deficiency causes the lysosomal storage disease sialidosis.^8–10^ The onset and severity of sialidosis symptoms varies across patients; those most affected are cognitively and neurodevelopmentally impaired.^8,9^

Sialidosis (*Neu1*^-/-^) mice develop a severe neurodegenerative disease with features of AD.^11^ In *Neu1*^-/-^ neurons, amyloid precursor protein (APP), a substrate of Neu1, remains sialylated and accumulates in lysosomes, where it is processed into a soluble fragment and a beta C-terminal fragment (CTF).^11^ This CTF is subsequently cleaved into amyloid beta (Aβ) peptides, which are released extracellularly through lysosomal exocytosis.^11^ Neu1 negatively regulates this physiological process by cleaving the sialic acids of lysosome-associated membrane protein 1 (LAMP1), shortening its half-life.^12^ Neu1 deficiency increases the number of PM-docked LAMP1^+^ lysosomes poised to fuse and release their contents extracellularly.^12^ Exacerbated lysosomal exocytosis compromises tissue integrity.^12^ In *Neu1*^-/-^ neurons, sialylated APP (Sia-APP) accumulates in lysosomes, thereby enhancing exocytosis of toxic Aβ peptides into the cerebrospinal fluid (CSF), which drives the development of an early amyloidosis phenotype.^11^

Another facet of AD pathology is microglia-mediated neuroinflammation.^13–15^ Microglia are the innate immune cells of the brain, and microglial subtypes are classified as responding, proinflammatory/neurotoxic, or anti-inflammatory/neuroprotective based on their microenvironment and intracellular status.^16^ In AD, microglia phagocytose extracellular debris and Aβ peptides, and they promote cellular responses to plaque deposition and neurodegeneration.^14^ In *Neu1*^-/-^ mice, increased exocytosis of Aβ-42, a potent microglia activator, most likely induces a microglia-mediated neuroinflammatory response resembling AD, which provides a unique platform to examine how Neu1 regulates microglial function in neurodegenerative diseases.

The microglial triggering receptor expressed on myeloid cells 2 (Trem2) mediates neuroinflammatory signaling in AD and contributes to microglia survival, phagocytosis, motility, proliferation, and cytokine production.^17–19^ Full-length Trem2 (Trem2-FL) requires interaction with the adaptor protein, DNAX-activating protein of 12 kDa (DAP12).^20^ High-avidity ligand binding to Trem2-FL–DAP12 complexes activates the receptor, triggering phosphorylation of the ITAM domain of DAP12, recruitment and phosphorylation of spleen tyrosine kinase (SYK), and intracellular signaling.^20^ SYK can induce protein kinase B (Akt) signaling, nuclear factor of activated T cells 1 (NFAT1) activation, phagocytosis, and nuclear factor kappa of B cells (NFκB) inhibition.^17,18,21–23^ The glycan composition of Trem2-FL is complex, including low molecular weight (∼30 kDa) species (Trem2-LMW), and higher molecular weight (35-60 kDa) species (Trem2-HMW).^24^ N-linked glycosylation of Trem2-FL affects its stability, trafficking to the PM, and post-activation signal transduction.^25^ However, the effects of sialylation on Trem2 remain elusive.

Trem2-dependent signaling is further compounded by the contribution of its cleavage products, an extracellular soluble domain (sTrem2) and a C-terminal fragment (Trem2-CTF), to inflammatory responses. Trem2-FL is cleaved by α-secretase, a disintegrin and metalloproteinase domain-containing protein 10 (ADAM10) or ADAM17, into Trem2-CTF and sTrem2.^26^ Trem2-CTF is further cleaved by γ-secretase into a β peptide and an intracellular domain.^27^ Trem2-CTF cleavage is suppressed by its interaction with DAP12, which stabilizes Trem2-CTF–DAP12 complexes and promotes NFκB inhibition, independent of Trem2-FL or SYK.^28^ The sequestration of DAP12 by Trem2-CTF also diminishes Trem2-FL signaling.^28^ Alternatively, sTrem2 suppresses apoptosis through Akt signaling, promotes cytokine production via the NFκB pathway, and triggers calcium (Ca^2+^) influx.^29,30^ Although both Trem2 proteolytic fragments partially mimic Trem2-FL functions, neither participates in Trem2-dependent phagocytosis.^31^

Here, we identified Trem2 as a substrate of Neu1. By cleaving the sialic acids on Trem2, Neu1 regulates its proteolytic processing, interaction with DAP12, and downstream signaling. In the absence of Neu1, microglia-mediated pro-inflammatory responses ensued within the hippocampus, where Sia–Trem2-FL accumulated at the PM and in endolysosomes. Sialylation of Trem2-HMW did not prevent its interaction with DAP12 but impeded Syk activation, thereby precluding Trem2-FL–dependent phagocytosis, Akt signaling, NFκB inhibition, and NFAT1 activation. Moreover, proteolytic processing of Sia–Trem2-HMW increased Trem2-CTF and sTrem2 levels at the PM and in lysosomes. This favored the formation of Trem2-CTF–DAP12 complexes, which promoted NFκB inhibition, and diminished the DAP12 pool that interacts with Trem2-HMW. Concurrently, Sia-sTrem2 did not induce the NFκB pathway, but constitutively activated Akt and NFAT1 signaling, thereby leading to the production of tumor necrosis factor alpha (TNFα) and C-C motif chemokine ligand 3 (CCL3) by Neu1-deficient microglia. Together these findings uncovered Neu1’s role in regulating Trem2 homeostasis and functions.

## Results

### *Neu1*^-/-^ microglia adopt a pro-inflammatory phenotype in hippocampi

We first characterized microglia in *Neu1*^-/-^ hippocampi. Compared to wild-type (WT) brains, *Neu1*^-/-^ brains contained more microglia at 5 months (Figure 1A) but not at 2 months (Figure S1A). This increase was evident in the CA3 region (Figure S1B), where amyloid deposits form. To quantify microglia cell count, we crossed WT and *Neu1*^-/-^ mice with CX3CR1^GFP^ mice, sorted cell populations using fluorescently activated cell sorting (FACS), and analyzed microglia by flow cytometry (Figure S1C). *Neu1*^-/-GFP^ hippocampi contained twice as many microglia as WT^GFP^ hippocampi did by 4 months (Figure 1B). Microglia shift their morphology from ramified to amoeboid upon activation. *Neu1*^-/-^ hippocampal microglia assumed an amoeboid morphology by 2 months (Figures 1C and S1A) and were significantly larger than WT microglia (Figures 1D and S1D), indicating their acquisition of an activated phenotype.

**Figure 1.**
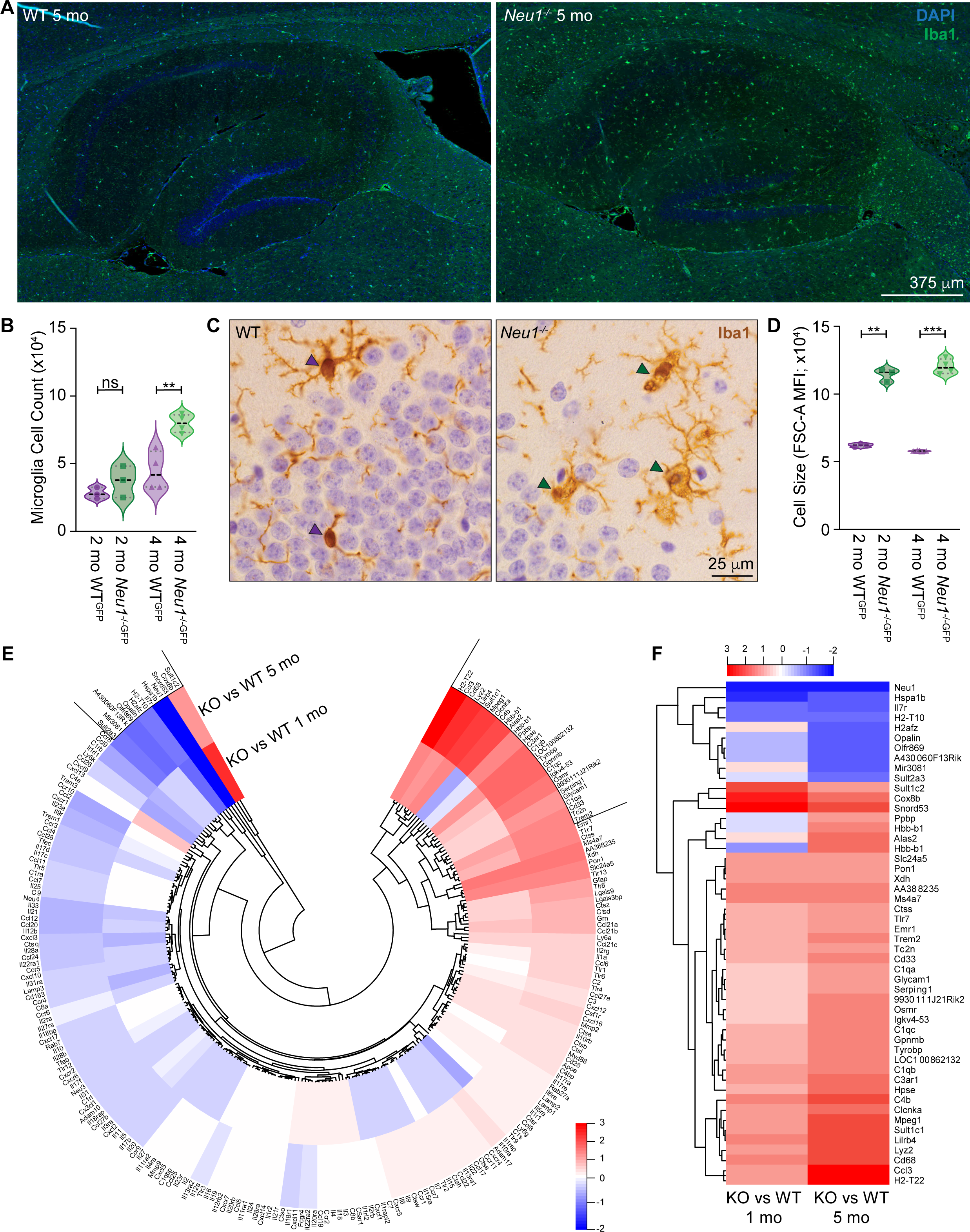
*Neu1*^-/-^ microglia are activated in hippocampi. (A) Immunofluorescence staining of microglia (Iba1; green) and nuclei (DAPI; blue) in 5-month-old WT and *Neu1*^-/-^ hippocampi. (B) Quantification of microglia (GFP^hi^) in 2- (n=3) and 4-month-old (n=4) WT^GFP^ and *Neu1*^-/-GFP^ hippocampi, as measured by FACS. (C) Immunohistochemical staining of microglia (Iba1; brown) in 2-month-old WT and *Neu1*^-/-^ hippocampi. Purple arrowheads indicate ramified (WT) morphology, and green arrowheads represent amoeboid (*Neu1*^-/-^) morphology. (D) Quantification of cell size by flow cytometry analysis of forward scatter area (FSC-A) of 2- (n=3) and 4-month-old (n=4) WT^GFP^ and *Neu1*^-/-GFP^ microglia. (E) Differential mRNA gene expression patterns in 1- and 5-month-old WT and *Neu1*^-/-^ hippocampi (n=3). Red: upregulation; blue: downregulation. Data represented as a polar heatmap with a dendrogram. (F) Top 40 upregulated (red) and top 10 downregulated (blue) genes from (E). Data represented as a heatmap with a dendrogram. Data represented as medians ± quartiles. ns: not significant, **p <0.01, ***p <0.001. See also Figure S1.

To characterize these cells, we analyzed RNA microarrays of 1- and 5-month-old WT and *Neu1*^-/-^ hippocampi (Figure 1E). The top 40 upregulated genes in *Neu1*^-/-^ samples belonged to the cytokine (*Ccl3*), phagocytic receptor (*Trem2*), and complement (*C1qa*) families (Figure 1F). Gene set enrichment analysis (GSEA) revealed several myeloid lineage–specific and/or pro-inflammatory pathways upregulated in *Neu1*^-/-^ hippocampi (Figure 2A-2C). GSEA also revealed the upregulation of specific pro-inflammatory cascades (Figure 2D-2F). Using qRT-PCR, we validated the expression of genes upregulated in the microarray and/or those encoding inflammatory cascade– associated proteins.

**Figure 2.**
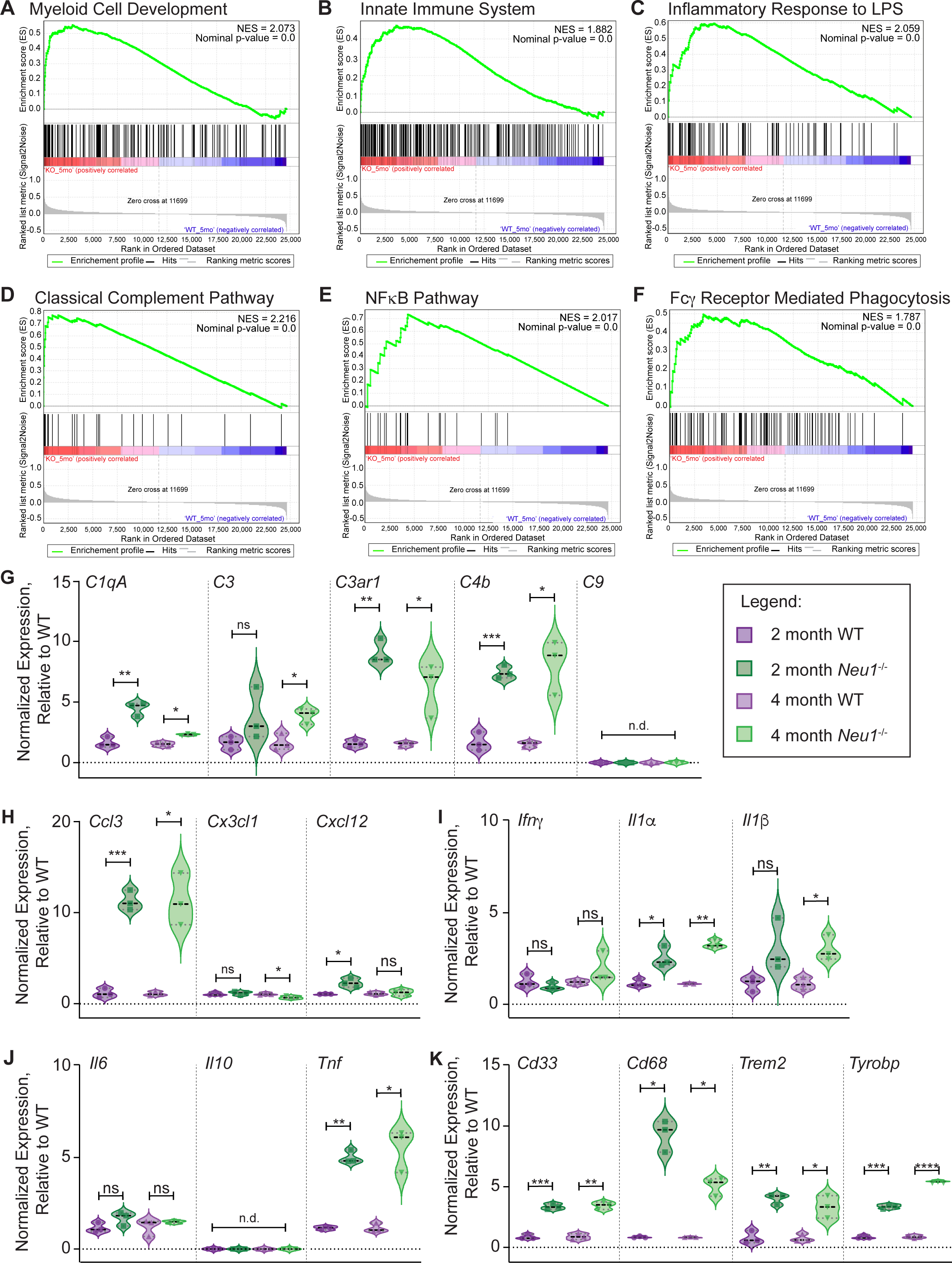
Genetic profiling of *Neu1*^-/-^ hippocampi. (A-F) Gene set enrichment analysis of differentially expressed genes (in Figure 1E) identified pathways involved in (A) myeloid cell development, (B) innate immune system,(C) inflammatory responses to lipopolysaccharide (LPS), (D) classical complement, (E) NFκB and (F) Fcγ receptor–mediated phagocytosis. False discovery rate (FDR) <0.25; NES: Nominal Enrichment Score. (G-K) The qRT-PCR analyses of gene expression in 2- and 4-month-old WT and *Neu1*^-/-^ hippocampi (n=3) for (G) *C1qa*, *C3*, *C3ar1*, *C4b*, and *C9*; (H) *Ccl3*, *Cx3cl1*, and *Cxcl12*; (I) *Ifnγ*, *Il1α*, and *Il1β*; (J) *Il6*, *Il10*, and *Tnf*; and (K) *Cd33*, *Cd68*, *Trem2*, and *Tyrobp*. Data represented as medians ± quartiles. n.d.= not detected, ns: not significant, *p <0.05, **p <0.01, ***p <0.001, ****p <0.0001.

Genes encoding the complement cascade–initiating factors, including complement component 1qA (*C1qa*) and *C3*, were upregulated in *Neu1*^-/-^ hippocampi (Figure 2G). Additionally, the expression of *C4b*, which encodes a required member of the C3-convertase, was increased (Figure 2G). The gene that encodes the C3a receptor 1 (C3ar1), which mediates downstream C3 signaling, was also upregulated in *Neu1*^-/-^ hippocampi (Figure 2G). However, *C9*, the terminal member of the classical complement cascade, was not detected in *Neu1*^-/-^ or WT samples at either age (Figure 2G), indicating that the cascade was not fully executed in *Neu1*^-/-^ mice.

We next analyzed cytokine and chemokine gene expression to define the *Neu1*^-/-^ hippocampal microenvironment as neurotoxic or neuroprotective. *Ccl3*, which encodes a neurotoxic chemokine, was upregulated in *Neu1*^-/-^ samples at 2 and 4 months (Figure 2H). In contrast, expression of C-X3-C motif chemokine ligand 1 (*Cx3cl1*), encoding a neuroprotective chemokine that promotes microglial survival, was not differentially expressed in 2-month-old *Neu1*^-/-^ hippocampi but was downregulated 40% in 4-month-old mice (Figure 2H). Likewise, expression of *Cxcl12*, which encodes a neuroprotective chemokine that supports neuronal survival, was upregulated in *Neu1*^-/-^ hippocampi at 2 months but normalized at 4 months (Figure 2H). We also detected upregulation of genes encoding the pro-inflammatory cytokines interleukin (*Il*)-*1α*, *Il-1β*, and *Tnf*, with *Tnf* being the most upregulated in *Neu1*^-/-^ hippocampi compared to WT (Figure 2I and 2J). The pro-inflammatory cytokine genes, interferon gamma (*Ifnγ)* and *Il-6* (Figure 2I and 2J), were not different between samples, and no expression of *Il-10*, an anti-inflammatory cytokine, was detected in *Neu1*^-/-^ or WT samples at any age (Figure 2J).

Upregulation of pro-inflammatory factors is associated with the modulation of phagocytosis genes. Therefore, we tested the expression of *Trem2, Cd68*, and *Cd33* and found upregulation of *Trem2* and *Cd68* in *Neu1*^-/-^ hippocampi at 2 and 4 months (Figure 2K). Similar patterns of upregulation were measured for *Tyrobp,* which encodes the Trem2 adaptor protein, DAP12, and *Cd33*, which encodes a sialic acid–binding receptor that modulates Trem2 signaling (Figure 2K). These results revealed a pro-inflammatory microenvironment in *Neu1*^-/-^ hippocampi and suggest that *Neu1*^-/-^ microglia have an increased phagocytic capacity.

### *Neu1*^-/-^ microglia secrete pro-inflammatory cytokines but have diminished phagocytosis

Lysosomal exocytosis, which is regulated by Neu1, is a mechanism by which immune cells release signaling molecules.^12,31^ To test the extent of exocytosis in *Neu1*^-/-^ hippocampal microglia, we assayed beta-hexosaminidase (β-Hex) activity in microglia culture media and quantified the pool of PM-localized Lamp1 (PM-Lamp1). *Neu1*^-/-^ microglia had higher β-Hex activity and PM-Lamp1 levels, establishing those cells as more exocytic than WT microglia (Figures 3A, 3B, and S2A). *Neu1*^-/-GFP^ microglia also released more TNFα and CCL3 than did WT^GFP^ controls (Figure 3C and 3D), confirming a microglia-mediated pro-inflammatory response in *Neu1*^-/-^ hippocampi. When lysosomal exocytosis was blocked in cultured microglia, CCL3 release was reduced, indicating that CCL3 is secreted by lysosomal exocytosis (Figure 3E).

**Figure 3.**
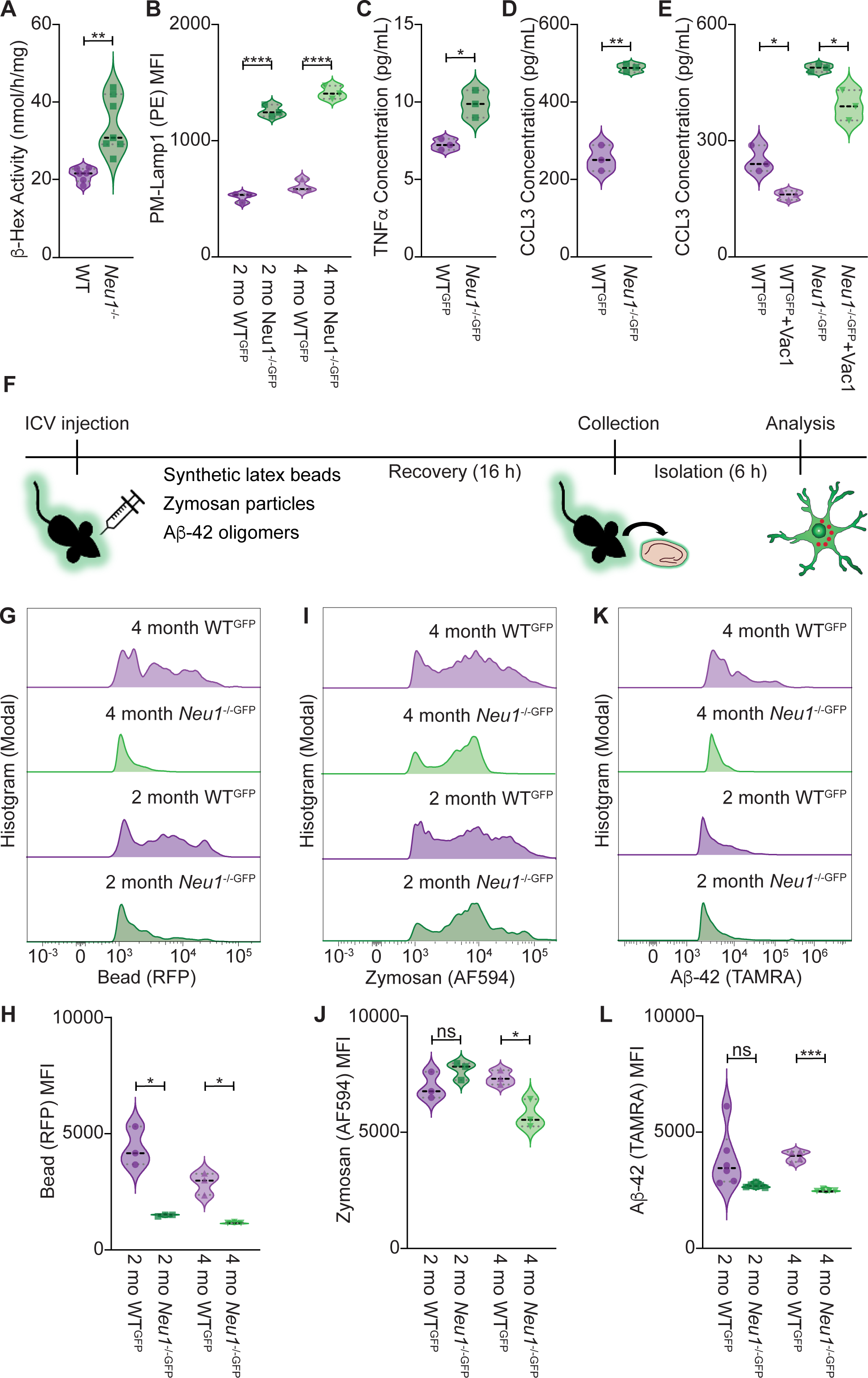
*Neu1*^-/-^ microglia secrete pro-inflammatory cytokines but have diminished phagocytosis. (A) Quantification of β-hexoxaminidase (β-Hex) activity in the media of WT (n=6) and *Neu1*^-/-^ (n=7) microglia cultures. (B) Quantification of PM-Lamp1 (PE) in 2- and 4-month-old WT^GFP^ and *Neu1*^-/-GFP^ microglia by FACS (n=3). (C and D) Concentration of (C) TNFα and (D) CCL3 in media from WT^GFP^ and *Neu1*^-/-GFP^ microglia cultures (n=3). (E) CCL3 concentrations in media from WT^GFP^ and *Neu1*^-/-GFP^ microglia cultures treated with 10 μM vacuolin-1 (Vac1) for 4 h (n=3). (F) Schematic of the *in vivo* phagocytosis assay procedures. (G-L) Representative plots and quantification of phagocytic capacity in 2- and 4-month-old WT^GFP^ and *Neu1*^-/-GFP^ microglia for (G-H) 1-μm red fluorescent latex beads (n=3), (I-J) AF594-labeled Zymosan A particles (n=3), and (K-L) TAMRA-labeled Aβ-42 oligomers (2 month: n=6; 4 month: n=4). Data represented as medians ± quartiles. *p <0.05, **p <0.01, ***p <0.001, ****p <0.0001. Abbreviations: ICV, intracerebral ventricular; MFI, mean fluorescence intensity; ns, not significant; RFP, red fluorescent protein See also Figure S2A.

We next tested multiple phagocytic pathways *in vivo* (Figure 3F). Mice were intracerebroventricularly injected with synthetic latex beads to induce immunoglobin G (IgG)-mediated phagocytosis, zymosan A particles to evoke phagocytosis through toll-like receptor 2/6, or Aβ-42 oligomers to stimulate phagocytosis and endocytosis. Unexpectedly, *Neu1*^-/-GFP^ microglia showed reduced phagocytosis of beads at 2- and 4-months, and of zymosan at 4 months (Figure 3G-3J). These cells also lost their ability to take up Aβ-42 oligomers by 2 months (Figure 3K and 3L), implicating Neu1 as a master regulator of microglia phagocytosis.

### Trem2 is differentially distributed and contributes to pro-inflammatory responses in *Neu1*^-/-^ microglia

We next investigated the subcellular distribution of Trem2 in *Neu1*^-/-^ hippocampal microglia. Only Trem2-HMW accumulated at the PM of *Neu1*^-/-GFP^ cells (Figures 4A-4C and S2B). Immunofluorescence staining showed a punctate intracellular distribution of Trem2-FL in *Neu1*^-/-GFP^ microglia (Figure 4D). Quantification of Trem2-FL^+^ objects revealed a higher intracellular volume in *Neu1*^-/-GFP^ microglia than in WT^GFP^ controls (Figure 4E). Object-intensity co-localization (OICL) measures revealed that Trem2-FL primarily localized in the Golgi of WT^GFP^ microglia but accumulated in endolysosomes of *Neu1*^-/-GFP^ microglia (Figures 4F and S2C-S2F). The largest pool of Trem2-FL in *Neu1*^-/-GFP^ microglia was in the lysosomes, and phagosome-localized Trem2-FL was seen only in *Neu1*^-/-GFP^ samples (Figures 4F, S2E, and S2F).

**Figure 4.**
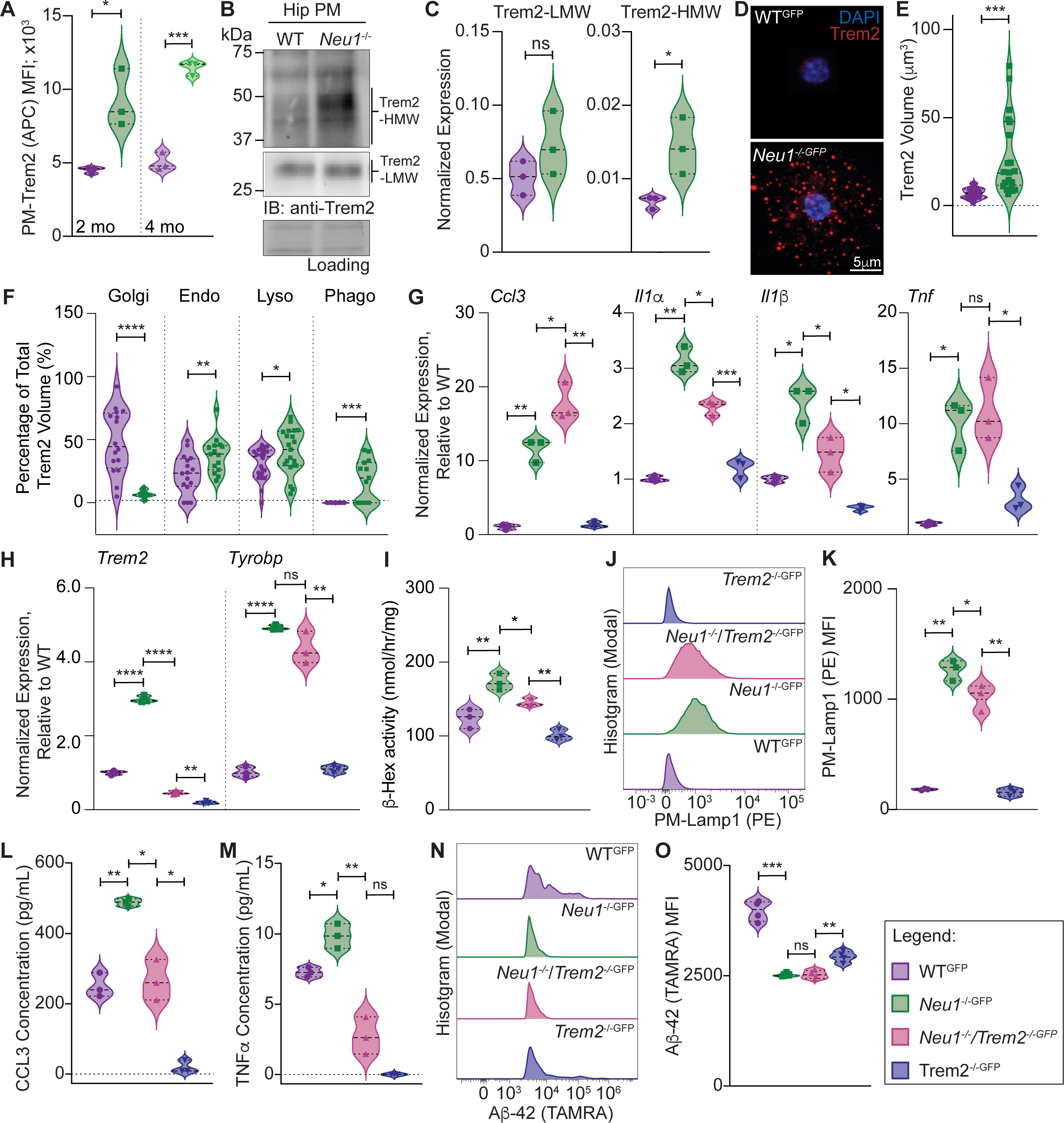
Trem2 is differentially distributed and contributes to pro-inflammatory responses in *Neu1*^-/-^ microglia. (A) Quantification of PM-Trem2-FL (APC) levels in 2- and 4-month-old WT^GFP^ and *Neu1*^-/-GFP^ microglia by FACS (n=3). (B-C) Representative immunoblot (B) and protein quantification (C; n=3) of Trem2-FL in the PM fractions of 4-month-old WT and *Neu1*^-/-^ hippocampi (Hip). Trem2-FL is separated into Trem2-HMW (right) and Trem2-LMW (left) species. (D-E) Representative immunofluorescence images (D) and volume quantification (E) of Trem2-FL^+^ (red) objects in 4-month WT^GFP^ (n=24) and *Neu1*^-/-GFP^ (n=21) microglia. DAPI (blue). (F) Quantification of Trem2^+^ object volume in various subcellular compartments of 4-month-old WT^GFP^ and *Neu1*^-/-GFP^ microglia. Golgi and Endo (early endosomes): WT^GFP^ (n=16), *Neu1*^-/-GFP^ (n=15); Lyso (lysosomes): WT^GFP^ (n=24), *Neu1*^-/-GFP^ (n=21); Phago (phagosomes): WT^GFP^ (n=22), *Neu1*^-/-GFP^ (n=15). (G-H) The qRT-PCR analyses of gene expression in 4-month-old WT^GFP^, *Neu1*^-/-GFP^, *Neu1*^-/-^/*Trem2*^-/-GFP^, and *Trem2*^-/-GFP^ hippocampi for (G) *Ccl3, Il1α*, *Il1β*, and *Tnf* and (H) *Trem2* and *Tyrobp* (n=3). (I) The β-Hex activity in media from WT^GFP^, *Neu1*^-/-GFP^, *Neu1*^-/-^/*Trem2*^-/-GFP^, and *Trem2*^-/-GFP^ microglia cultures (n=3). (J) Quantification of PM-Lamp1 levels in WT^GFP^, *Neu1*^-/-GFP^, *Neu1*^-/-^/*Trem2*^-/-GFP^, and *Trem2*^-/-GFP^ microglia by FACS (n=3). (K) Representative plot for PM-Lamp1 expressed in 4-month-old WT^GFP^, *Neu1*^-/-GFP^, *Neu1*^-/-^/*Trem2*^-/-GFP^, and *Trem2*^-/-GFP^ microglia. (L-M) Concentrations of (L) CCL3 and (M) TNFα in the media of WT^GFP^, *Neu1*^-/-GFP^, *Neu1*^-/-^/*Trem2*^-/-GFP^, and *Trem2*^-/-GFP^ microglia (n=3). (N-O) Representative plot (N) and quantification (O) of phagocytic capacity for TAMRA-labeled Aβ-42 oligomers in WT^GFP^, *Neu1*^-/-GFP^, *Neu1*^-/-^/*Trem2*^-/-GFP^, and *Trem2*^-/-GFP^ microglia (n=4). Data represented as the medians ± quartiles; ns: not significant, *p <0.05, **p <0.01, ***p <0.001, ****p <0.0001. See also Figure S2B-S2K.

To establish a hierarchical relationship between Neu1 and Trem2 in microglia, we crossed *Neu1*^-/-GFP^ mice with *Trem2*^-/-^ mice and compared the expression of inflammatory genes in WT^GFP^, *Neu1*^-/-GFP^, *Neu1*^-/-^/*Trem2*^-/-GFP^, and *Trem2*^-/-GFP^ hippocampi. Expression of *Il-1α* and *Il-1β* was lower, but that of *Ccl3* was higher in *Neu1*^-/-^/*Trem2*^-/-GFP^ hippocampi than in *Neu1*^-/-GFP^ hippocampi (Figure 4G). *Tnf*, *Ifnγ*, *Il-6*, *Il-10*, *Cx3cl1*, and *Cxcl12* remained unchanged (Figures 4G, S2G, and S2H). Of the genes associated with phagocytosis, only *Cd68* expression was higher in *Neu1*^-/-^/*Trem2*^-/-GFP^ hippocampi; *Cd33* and *Tyrobp* remained unchanged (Figures 4H and S2I). As expected, *Trem2* expression was lower in Trem2-deficient hippocampi (Figure 4H). Complement gene expression was also unchanged (Figure S2J). *Neu1*^-/-^/*Trem2*^-/-GFP^ hippocampi genetically resembled *Neu1*^-/-GFP^ hippocampi more than *Trem2*^-/-GFP^ hippocampi, indicating that Neu1 is hierarchically above Trem2 (Figures 4G, 4H, and S2G-S2J). The GFP transgene did not affect microglia-mediated responses; WT^GFP^ and *Neu1*^-/-GFP^ genetic profiles were comparable to WT and *Neu1*^-/-^ profiles (Figures 2G-2K, 4G, 4H, and S2G-S2J).

We next conducted functional studies of *Neu1*^-/-^/*Trem2*^-/-GFP^ microglia to elucidate the contribution of Trem2 to *Neu1*^-/-^ microglia dysfunction. We measured β-Hex activity in culture media and PM-Lamp1 levels in WT^GFP^, *Neu1*^-/-GFP^, *Neu1*^-/-^/*Trem2*^-/-GFP^, and *Trem2*^-/-GFP^ microglia. Both β-Hex activity and PM-Lamp1 levels were reduced in *Neu1*^-/-^/*Trem2*^-/-GFP^ cells compared to *Neu1*^-/-GFP^ cells (Figure 4I-4K). Moreover, *Neu1*^-/-^/*Trem2*^-/-GFP^ microglia released less CCL3 and TNFα than did *Neu1*^-/-GFP^ cells (Figure 4L and 4M). These results suggest that Trem2 contributes to lysosomal exocytosis and cytokine release from *Neu1*^-/-GFP^ cells.

Using our *in vivo* approach, we determined that the uptake of Aβ-42 oligomers was similar between *Neu1*^-/-^/*Trem2*^-/-GFP^ and *Neu1*^-/-GFP^ microglia (Figure 4N and 4O). *Trem2*^-/-GFP^ microglia maintained the ability to phagocytose Aβ-42 oligomers, suggesting that Neu1 is upstream of Trem2 in Trem2-dependent phagocytosis (Figure 4N and 4O). These results suggest that Trem2 signaling contributes to the pro-inflammatory phenotype of *Neu1*^-/-^ microglia.

### Trem2 sialylation promotes its proteolytic processing and accumulation of its cleaved products

We next compared the levels of Sia–Trem2-FL between WT^GFP^ and *Neu1*^-/-GFP^ primary microglia. Using OICL of Sambucus nigra (SNA) and Trem2, we found a higher volume of Sia–Trem2-FL in *Neu1*^-/-GFP^ microglia than in WT^GFP^ cells (Figures 5A and S3A). Immunoblot analyses of microglia lysates confirmed increased Trem2-HMW levels and accumulated Trem2-CTF in *Neu1*^-/-GFP^ samples (Figure 5B). Concomitantly, sTrem2 was increased in the soluble fraction of *Neu1*^-/-^ hippocampi, *Neu1*^-/-^ CSF, and *Neu1*^-/-GFP^ microglia culture medium (Figure 5C).

**Figure 5.**
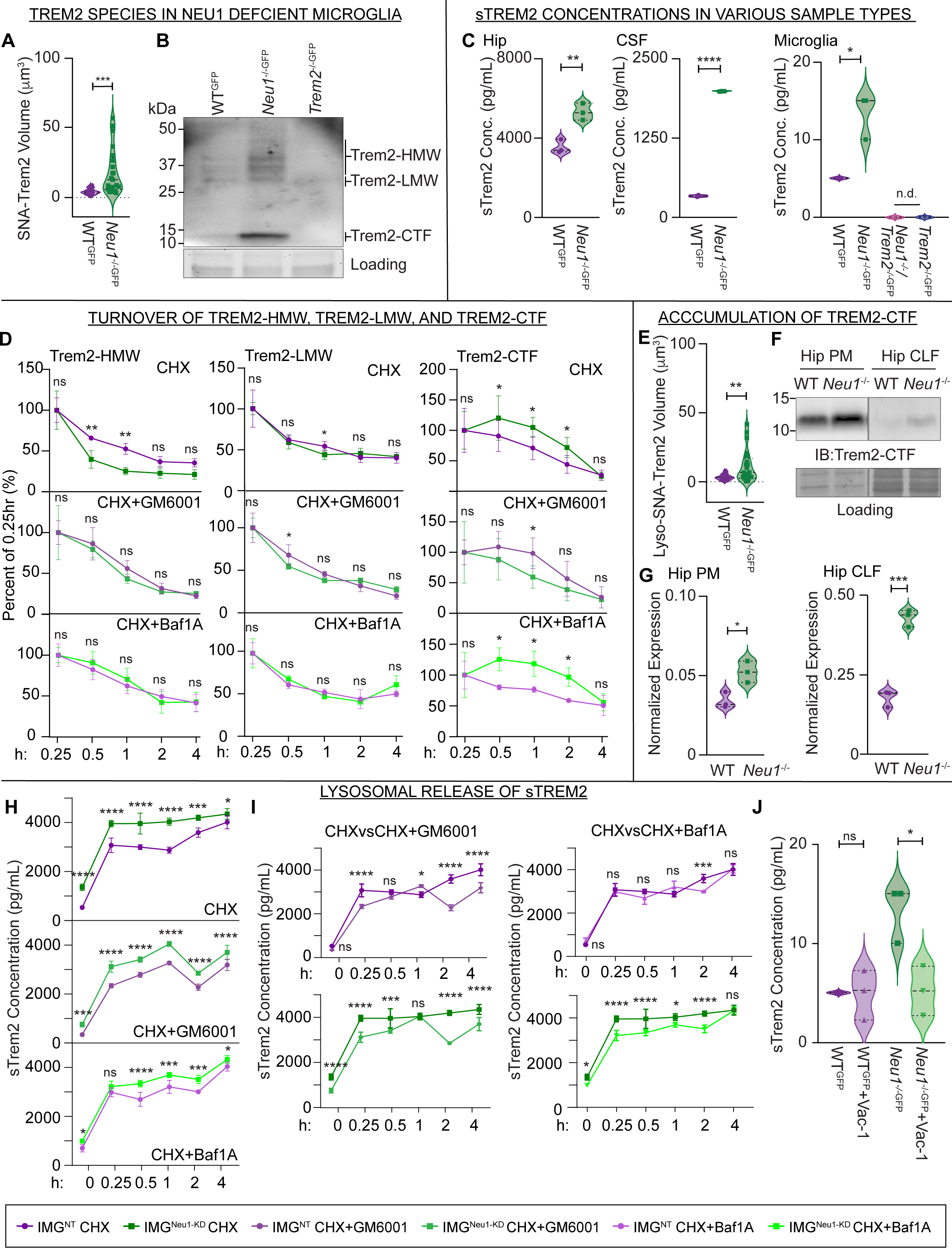
Trem2 sialylation promotes the proteolytic processing and accumulation of its cleavage products. (A) Quantification of Trem2^+^/SNA^+^ object volume in 4-month-old WT^GFP^ (n=24) and *Neu1*^-/-GFP^ (n=21) microglia. (B) Immunoblot of total Trem2 (i.e., Trem2-HMW, Trem2-LMW, and Trem2-CTF) in 4-month-old WT^GFP^, *Neu1*^-/-GFP^, and *Trem2*^-/-GFP^ microglia cell lysates. (C) Concentration of sTrem2 in (left) hippocampal (Hip) soluble fractions, (middle) CSF from 4-month-old WT^GFP^ and *Neu1*^-/-GFP^ mice, and (right) media from WT^GFP^, *Neu1*^-/-GFP^, *Neu1*^-/-^/*Trem2*^-/-GFP^, and *Trem2*^-/-GFP^ microglia cultures (n=3). (D) Quantification of (left) Trem2-HMW, (middle) Trem2-LMW, and (right) Trem2-CTF in IMG^NT^ and IMG^Neu1-KD^ cells treated with CHX (n=5), CHX+GM6001 (n=4), or CHX+Baf1A (n=4) for 0.25, 0.5, 1, 2, or 4 h; data represented as a percentage of the concentration at 0.25 h. (E) Quantification of Trem2^+^/SNA^+^ object volume in lysosomes of 4-month-old WT^GFP^ (n=24) and *Neu1*^-/-GFP^ (n=21) microglia. (F-G) Representative immunoblots (F) and quantification of Trem2-CTF in the (G, left) PM and (G, right) lysosomal (CLF) fractions of 4-month-old WT and *Neu1*^-/-^ hippocampi (n=3). (H) Concentration of sTrem2 in the media of IMG^NT^ and IMG^Neu1-KD^ cells in culture treated with CHX, CHX+GM6001, or CHX+Baf1A (n=4) for 0, 0.25, 0.5, 1, 2, or 4 h. (I) Comparison of sTrem2 concentrations between (left) CHX and CHX+GM6001 and (right) CHX and CHX+Baf1A treatments of IMG^NT^ (top) and IMG^Neu1-KD^ (bottom) cells at 0, 0.25, 0.5, 1, 2, and 4 h (n=4). (J) The concentration of sTrem2 in media from WT^GFP^ and *Neu1*^-/-GFP^ microglia cultures treated with 10 μM vacuolin-1 (Vac1) for 4 h (n=3). Data represented as mean ± SEMs; ns: not significant, n.d.= not detected, *p <0.05, **p <0.01, ***p <0.001, ****p <0.0001. See also Figures S3 and S4.

To elucidate the contribution of metalloproteinases and lysosomes to Trem2 processing, we performed cycloheximide (CHX)-turnover assays in IMG^NT^ and IMG^Neu1-KD^ cells; results are summarized in Table S1. For Trem2-FL, IMG^Neu1-KD^ cells treated with only CHX showed enhanced processing of Trem2-HMW but not Trem2-LMW, compared to that in IMG^NT^ cells (Figure 5D). Conversely, cells treated with CHX, with or without GM6001 or bafilomycin A1 (BafA1), showed no differences in Trem2-HMW or Trem2-LMW processing (Figure 5D). The nonsignificant point in Trem2-HMW levels between IMG^Neu1-KD^ and IMG^NT^ cells was reached at 0.5 h in CHX-treated cells, at 1 h in CHX+GM6001-treated cells, and at 2 h in CHX+Baf1A-treated cells (Figure S3B and S3C). This result indicated that both metalloproteinases and lysosomal degradation exacerbate the processing of Trem2-HMW in IMG^Neu1-KD^ cells. We found no major differences in Trem2-LMW levels between IMG^Neu1-KD^ and IMG^NT^ samples under any treatment (Figure S3B and S3D). Furthermore, Sia–Trem2-FL volume increased in lysosomes of *Neu1*^-/-GFP^ microglia compared to that in WT^GFP^ cells (Figures 5E and S4A). Thus, Neu1 deficiency drives preferential proteolysis of Sia–Trem2-HMW at the PM and in lysosomes to increase Trem2-CTF and sTrem2 levels.

To ascertain where Trem2-CTF accumulated, we isolated the PM and crude lysosomal fractions from *Neu1*^-/-GFP^ and WT^GFP^ hippocampi; Trem2-CTF accumulated at both sites in *Neu1*^-/-GFP^ cells (Figure 5F and 5G). We used CHX assays to monitor Trem2-CTF processing in Neu1-deficient microglia. When we treated IMG^Neu1-KD^ and IMG^NT^ cells with only CHX, Trem2-CTF degradation was slower in IMG^Neu1-KD^ cells than in IMG^NT^ cells, resulting in higher levels of Trem2-CTF until the 4-h time point (Figures 5D, S4B, and S4C). Treating cells with CHX+GM6001 abolished the differences in Trem2-CTF degradation, but Trem2-CTF levels remained higher in IMG^Neu1-KD^ cells until the 1-h time point (Figures 5D, S4B, and S4C). Therefore, Neu1-deficient microglia cannot compensate for enhanced cleavage of Trem2-HMW by metalloproteinases. Conversely, CHX+Baf1A treatment inhibited Trem2-CTF processing by slowing Trem2-CTF turnover, which resulted in higher levels at all time points in IMG^Neu1-KD^ cells (Figures 5D, S4B, and S4C). Thus, the lysosome is the primary site of Trem2-CTF degradation in IMG^Neu1-KD^ cells.

We next measured the concentration of sTrem2 released into the medium of cells treated with CHX alone, CHX+GM6001, or CHX+BafA1. The sTrem2 levels were consistently highest in IMG^Neu1-KD^ medium (Figure 5H). Compared to the effect of CHX alone, the differences between cell lines were smaller but still significant after CHX+GM6001 or CHX+BafA1 treatment, implicating both the PM and lysosomes as sites of sTrem2 production. However, when we compared the sTrem2 levels across treatment groups of the same cell line, only IMG^Neu1-KD^ cells showed differences between CHX-only and CHX+BafA1 treatments, whereas both cell lines showed differences between CHX-only and CHX+GM6001 treatments (Figure 5I). Thus, only IMG^Neu1-KD^ cells produced a pool of sTrem2 in lysosomes that enhanced the extracellular levels of sTrem2. Next, we inhibited lysosomal exocytosis in WT^GFP^ and *Neu1*^-/-GFP^ microglia; only *Neu1*^-/-GFP^ microglia released lower levels of sTrem2 (Figure 5J). These results confirmed that the difference in sTrem2 levels between *Neu1*^-/-GFP^ and WT^GFP^ microglia was due to its excessive release via lysosomal exocytosis.

### Sialylation of Trem2-HMW influences its association with DAP12

To investigate the association of Trem2 with DAP12 in Neu1-deficient microglia, we established an IMG cell line with dual downregulation of *Neu1* and *Trem2* (IMG^N1/T2-dKD^). IMG^NT^, IMG^Neu1-KD^, and IMG^N1/T2-dKD^ cells were stimulated with CSF collected from 4-month-old WT (homeostatic) or *Neu1^-/-^* (pro-inflammatory) mice; results are summarized in Table S2. When stimulation shifted from homeostatic to pro-inflammatory, Trem2-HMW levels increased in IMG^Neu1-KD^ cells but decreased in IMG^NT^ cells; Trem2-LMW levels decreased in both lines (Figures 6A and S5A). IMG^Neu1-KD^ cells consistently maintained higher levels of both Trem2-FL species compared to those in IMG^NT^ cells (Figure S5B and S5C). Therefore, Neu1 enzymatically acts only on Trem2-HMW but indirectly regulates all Trem2-FL levels across microglia cellular states.

**Figure 6.**
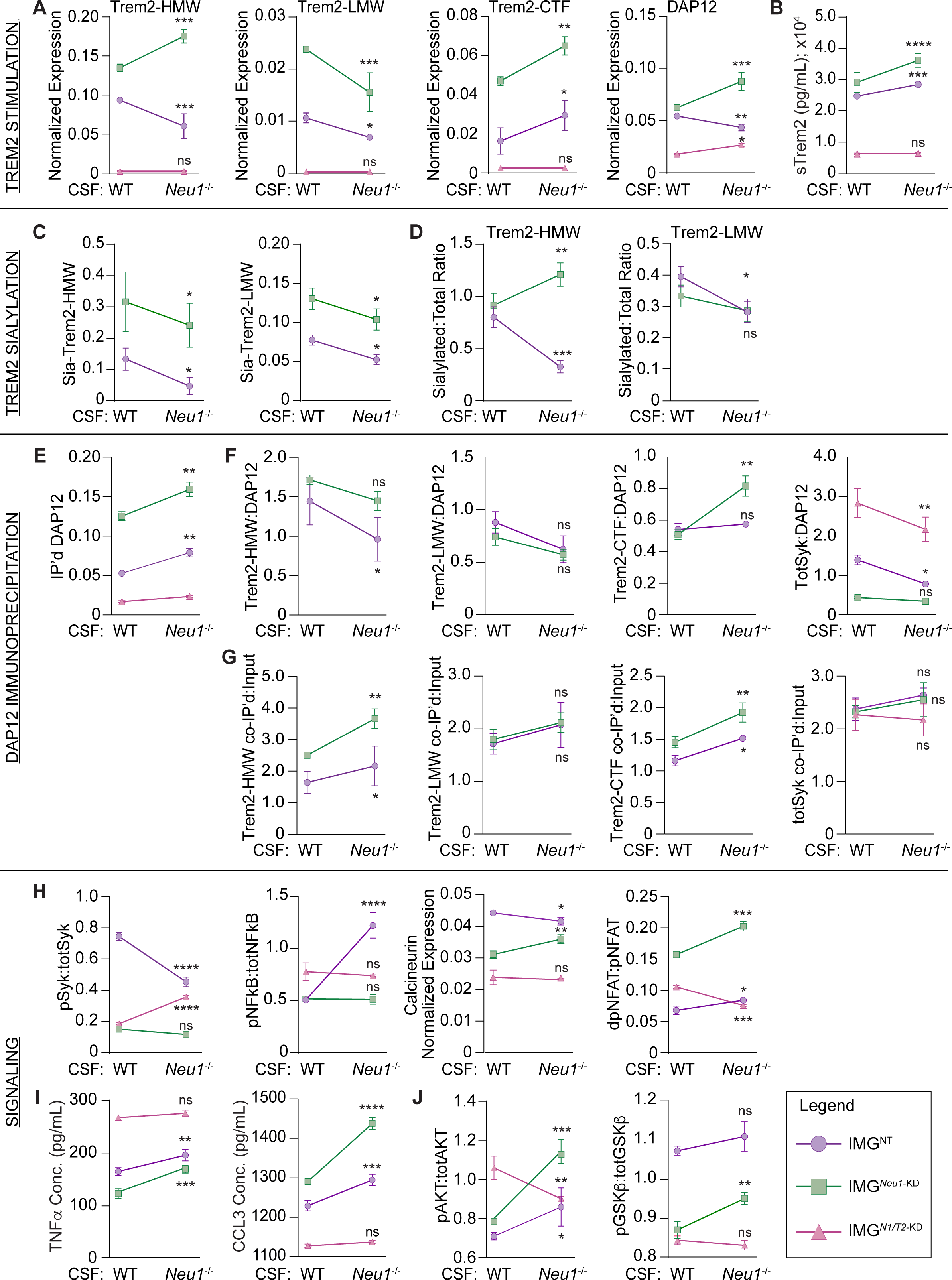
Sialylation of Trem2-HMW influences its association with DAP12 and alters its downstream signaling. (A) Normalized expression of (left) Trem2-HMW, (middle-left) Trem2-LMW, (middle-right) Trem2-CTF, and (right) DAP12 in stimulated IMG^NT^, IMG^Neu1-KD^, and IMG^N1/T2-dKD^ cells. (B) The concentration of sTrem2 in media from stimulated IMG^NT^, IMG^Neu1-KD^, and IMG^N1/T2-dKD^ cells. (C) Normalized expression of (left) Sia–Trem2-HMW and (right) Sia–Trem2-LMW in stimulated IMG^NT^ and IMG^Neu1-KD^ cells. (D) The ratio of sialylated (left) Trem2-HMW and (right) Trem2-LMW to total Trem2 in stimulated IMG^NT^ and IMG^Neu1-KD^ cells. (E) Normalized expression of immunoprecipitated (IP’d) DAP12 from stimulated IMG^NT^, IMG^Neu1-KD^, and IMG^N1/T2-dKD^ cells. (F-G) The ratios of co-immunoprecipitated (left) Trem2-HMW, (middle-left) Trem2-LMW, (middle-right) Trem2-CTF, and (right) Syk to immunoprecipitated DAP12 and (G, same order) to their corresponding input from stimulated IMG^NT^ and IMG^Neu1-KD^ cells. IMG^N1/T2-dKD^ cells were included in Syk calculations. (H) Signaling molecule activation, represented as the ratio of (left) pSyk^y525/526^ and (middle-left) pNFκB^S536^ to their total protein levels, (middle-right) normalized expression of calcineurin, and (right) the ratio of dpNFAT1 to pNFAT1 in stimulated IMG^NT^, IMG^Neu1-KD^, and IMG^N1/T2-dKD^ cells. (I) Concentrations of (left) TNFα and (right) CCL3 in media from stimulated IMG^NT^, IMG^Neu1-KD^, and IMG^N1/T2-dKD^ cells. (J) Signaling molecule activation, represented as the ratio of (left) pAkt^S473^ and (right) pGSK3β^S9^ to their total protein levels in stimulated IMG^NT^, IMG^Neu1-KD^, and IMG^N1/T2-dKD^ cells. (A-J) All samples were stimulated for 24 h with CSF collected from 4-month-old WT or *Neu1*^-/-^ mice (n=3 for all panels). Data represented as the mean ± SEM. ns: not significant, *p <0.05, **p <0.01, ***p <0.001, ****p <0.0001. See also Figures S5 and S6.

DAP12 levels increased in IMG^Neu1-KD^ but decreased in IMG^NT^ cells when stimulation shifted from homeostatic to pro-inflammatory (Figures 6A and S5A). DAP12 levels also increased in IMG^N1/T2-dKD^ cells, suggesting that these increases were a direct consequence of Neu1 downregulation (Figures 6A and S5A). Under homeostatic conditions, DAP12 levels were comparable between IMG^NT^ cells and IMG^Neu1-KD^ cells, but under pro-inflammatory conditions, they increased in IMG^Neu1-KD^ cells, suggesting that Neu1 influences DAP12 expression after microglial activation (Figure S5D). DAP12 levels were consistently lowest in IMG^N1/T2-dKD^ cells, indicating that DAP12 expression is always Trem2 dependent (Figure S5D).

Next, we found that Trem2-CTF levels and sTrem2 release were increased in IMG^Neu1-KD^ and IMG^NT^ cells, when the stimulus was shifted, but remained the highest in IMG^Neu1-KD^ cells under both conditions (Figures 6A, 6B, S5A, S5E, and S5F). As expected, Trem2-CTF and sTrem2 were not detected above background levels in IMG^N1/T2-dKD^ cells (Figures 6A, 6B, S5A, S5E, and S5F). These data indicate that Trem2 processing is promoted in pro-inflammatory microenvironments, leading to excessive production of Trem2-CTF and sTrem2, particularly when Neu1 is downregulated.

To determine whether Trem2-FL is a substrate of Neu1, we used biotinylated SNA to immunoprecipitate total sialylated proteins from stimulated IMG^NT^ and IMG^Neu1-KD^ cell lysates and performed immunoblot analyses for Trem2-FL; results are summarized in Table S3. Sia–Trem2-FL levels decreased in both cell lines when stimulation shifted to pro-inflammatory but remained higher in IMG^Neu1-KD^ cells under all conditions (Figures 6C and S5G-S5I). To assess changes in Trem2-FL sialylation after stimulation, we calculated the ratio of Sia–Trem2-HMW and Sia–Trem2-LMW to their total levels (sialylated:total). Sialylated:total Trem2-HMW did not differ between IMG^NT^ and IMG^Neu1-KD^ cells under homeostatic conditions but was higher in IMG^Neu1-KD^ cells under pro-inflammatory conditions (Figure S5J). This change was due to opposing responses of IMG^NT^ and IMG^Neu1-KD^ cells to the pro-inflammatory stimulus because the sialylated:total Trem2-HMW ratio increased in IMG^Neu1-KD^ cells but decreased in IMG^NT^ cells (Figure 6D). Conversely, we detected no differences in sialylated:total Trem2-LMW between IMG^NT^ and IMG^Neu1-KD^ cells under any conditions (Figures 6D and S5K). Thus, under pro-inflammatory conditions, Trem2-HMW undergoes Neu1-mediated de-sialylation, classifying it as a Neu1 substrate.

To test whether Trem2-FL sialylation influences its interaction with DAP12, we co-immunoprecipitated DAP12 with Trem2 species or Syk from stimulated IMG^NT^, IMG^Neu1-KD^, and IMG^N1/T2-dKD^ cells (Figure S5L); results are summarized in Table S4. Increased amounts of DAP12 were immunoprecipitated from all cell lines under pro-inflammatory conditions; the highest amounts were always from IMG^Neu1-KD^ cells (Figures 6E and S5M). First, to measure the proportion of DAP12 occupied by each protein (Trem2-HMW, Trem2-LMW, Trem2-CTF, or Syk), we calculated the ratio between the amount of each protein co-immunoprecipitated with DAP12 and the amount of immunoprecipitated DAP12 (protein:DAP12). Second, to evaluate the portion of each protein that was complexed with DAP12, we calculated the ratio between the amount of each protein co-immunoprecipitated and its own input value (protein co-IP’d:input). No Trem2 species were co-immunoprecipitated in IMG^N1/T2-dKD^ cells (Figure S5L). When stimulation was shifted to pro-inflammatory, Trem2-HMW:DAP12 decreased in IMG^NT^ cells but did not change in IMG^Neu1-KD^ cells (Figure 6F), indicating that this microenvironment antagonizes DAP12’s association with Trem2-HMW in IMG^NT^ cells but not in IMG^Neu1-KD^ cells. There was no difference in Trem2-HMW:DAP12 between IMG^NT^ and IMG^Neu1-KD^ cells under homeostatic conditions. However, this ratio was higher in IMG^Neu1-KD^ cells under pro-inflammatory stimulation (Figure S5N), demonstrating that DAP12 remains in complex with Trem2-HMW only in IMG^Neu1-KD^ cells. The Trem2-HMW co-IP’d:input ratio was consistently higher in IMG^Neu1-KD^ cells (Figure S5O), demonstrating that Trem2-HMW preferentially complexes with DAP12 when Neu1 is downregulated. When stimulation was shifted, the Trem2-HMW co-IP’d:input ratio increased in IMG^Neu1-KD^ cells but remained unchanged in IMG^NT^ cells (Figure 6G). This result suggests that changes in microglial cellular state affect only the association of Sia–Trem2-HMW with DAP12. In contrast, the Trem2-LMW:DAP12 and Trem2-LMW co-IP’d:input ratios showed no differences between IMG^NT^ and IMG^Neu1-KD^ cells under any conditions (Figures 6F, 6G, S5P, and S5Q), again demonstrating that Neu1 acts preferentially on Trem2-HMW. Overall, Neu1-mediated de-sialylation of Trem2-HMW regulated its complex formation with DAP12.

We next analyzed interactions between Trem2-CTF and DAP12. When stimulation was shifted, Trem2-CTF:DAP12 did not change in IMG^NT^ cells but almost doubled in IMG^Neu1-KD^ cells (Figures 6F and S6A). Shifting the stimulus caused the Trem2-CTF co-IP’d:input to increase in both IMG^NT^ and IMG^Neu1-KD^ cells, but this ratio was always higher in IMG^Neu1-KD^ cells (Figures 6G and S6B). Thus, retaining sialic acids on Trem2-HMW favored prolonged Trem2-CTF–DAP12 association under pro-inflammatory conditions.

Lastly, we analyzed how altering the formation of the Trem2–DAP12 complex influenced its subsequent interaction with Syk. When stimulation was shifted, the total Syk (totSyk):DAP12 ratio decreased in both IMG^NT^ cells and IMG^N1T2-dKD^ cells but remained unchanged in IMG^Neu1-KD^ cells (Figure 6F). With either stimulus, IMG^N1/T2-dKD^ cells maintained the highest totSyk:DAP12 ratio, and IMG^Neu1-KD^ cells maintained the lowest (Figure S6C). The calculation of this ratio was skewed by the differences in the amount of DAP12 immunoprecipitated from IMG^Neu1-KD^ and IMG^N1/T2-dKD^ cells. Therefore, Neu1 modulated the amount of DAP12 available to interact with Syk in a Trem2-dependent manner. Syk co-IP’d:input did not differ across cell lines or stimulation conditions (Figures 6G and S6D), indicating that neither Neu1 nor Trem2 affected Syk’s interaction with DAP12. Thus, Neu1-dependent de-sialylation of Sia–Trem2-HMW controls its interaction with DAP12, regulates the recruitment and retention of Syk, and influences the stability of Trem2-CTF–DAP12 complexes.

### Neu1 downregulation alters microglial signaling downstream of Trem2 activation

We analyzed the activation of intracellular molecules that participate in Trem2-FL, Trem2-CTF, and sTrem2 signaling;^17,18,28–30^ results are summarized in Table S2. First, we calculated levels of activated Syk, represented as the ratio of phosphorylated Syk at Y525/526 to total Syk (pSyk:totSyk), in stimulated IMG^NT^, IMG^Neu1-KD^, and IMG^N1/T2-dKD^ cells. IMG^NT^ cells maintained the highest levels of activated Syk, and IMG^Neu1-KD^ cells had the lowest, under all conditions (Figure S6E); therefore, Neu1 affected Syk phosphorylation downstream of Trem2 activation. When we shifted the stimulus, Syk activation decreased in IMG^NT^ cells but not in IMG^Neu1-KD^ cells (Figure 6H), demonstrating that a pro-inflammatory microenvironment diminishes Syk activation. The recovery of Syk activation in IMG^N1/T2-dKD^ cells under pro-inflammatory conditions (Figure 6H) revealed an unknown mechanism of Syk activation that occurs only when Neu1 and Trem2 are downregulated.

Because Trem2-FL–DAP12–Syk and Trem2-CTF–DAP12 signaling antagonize NFκB,^28^ we measured NFκB activation expressed as the ratio of phosphorylated NFκB p65 at S536 to total NFκB (pNFκB:totNFκB). NFκB was robustly activated in IMG^NT^ cells under pro-inflammatory stimulation, paralleling reductions in Syk activation, but under homeostatic conditions NFκB activation was comparable between IMG^NT^ cells and IMG^Neu1-KD^ cells (Figure S6F). This result confirmed that Neu1 influences NFκB activation. When stimulation shifted, there was no change in NFκB activation between IMG^Neu1-KD^ and IMG^N1/T2-dKD^ cells (Figure 6H). However, IMG^N1/T2-dKD^ cells consistently maintained higher levels of activated NFκB, indicating that the diminished NFκB activation in IMG^Neu1-KD^ cells was Trem2 dependent. We attributed this to stabilized Trem2-CTF–DAP12 complexes, given that Trem2-HMW–DAP12–Syk complexes did not transduce signal to subsequently antagonize NFκB. The lack of NFκB activation in IMG^Neu1-KD^ cells further suggested that sTrem2, which is highest in IMG^Neu1-KD^ culture medium, could not activate the pathway, most likely due to its sialylation status.

Despite impaired NFκB signaling in IMG^Neu1-KD^ cells, *Neu1^-/-^* cells still released TNFα and CCL3 (Figure 3C and 3D). Therefore, we measured NFAT1 activation, which induces the expression of these cytokines.^32,33^ Trem2-FL and sTrem2 may activate NFAT1, and Trem2-CTF may indirectly inhibit it by sequestering DAP12 in complex.^23,30^ NFAT1 is activated downstream of Ca^2+^-dependent calcineurin phosphatase activity. Therefore, we examined calcineurin levels in stimulated cells and found that IMG^NT^ cells unfailingly maintained the highest calcineurin levels, followed by IMG^Neu1-KD^ cells and IMG^N1/T2-dKD^ cells (Figure S6G). This result suggests that Neu1 and Trem2 are required to maintain normal calcineurin levels. However, when stimulation was shifted, calcineurin levels increased only in IMG^Neu1-KD^ cells, while they decreased in IMG^NT^ cells and did not change in IMG^N1/T2-dKD^ cells (Figure 6H). Thus, Neu1 regulates calcineurin in a Trem2-dependent manner.

We next measured NFAT1 activation as the ratio of dephosphorylated (active) to phosphorylated (inactive) protein (dpNFAT1:pNFAT1). IMG^Neu1-KD^ cells had the highest levels of activated NFAT1 (Figure S6H). Under homeostatic conditions, NFAT1 activation was higher in IMG^N1/T2-dKD^ cells than in IMG^NT^ cells, but under pro-inflammatory stimulation, the NFAT1 activation was comparable (Figure S6H). Thus, downregulation of Neu1 promotes constitutive NFAT1 activation in a Trem2-dependent manner. When stimulation was shifted, NFAT1 activation increased in IMG^NT^ cells and IMG^Neu1-KD^ cells but decreased in IMG^N1/T2-dKD^ cells (Figure 6H). Therefore, one or more Trem2 species are required for NFAT1 activation when microglial state changes; in IMG^Neu1-KD^ cells, we attributed NFAT1 activation to sTrem2, because Trem2-FL–DAP12–Syk does not transduce signal.

As a readout of these signaling events, we measured TNFα and CCL3 levels in stimulated IMG^NT^, IMG^Neu1-KD^, and IMG^N1/T2-dKD^ culture media. With either stimulus, IMG^NT^ cells released more TNFα but less CCL3 than did IMG^Neu1-KD^ cells; IMG^N1/T2-dKD^ cells released the most TNFα but the least CCL3 overall (Figure S6I and S6J). Thus, both Neu1 and Trem2 modulate the release of these cytokines. When stimulation was shifted, IMG^NT^ cells and IMG^Neu1-KD^ cells released more TNFα and CCL3 (Figure 6I). CCL3 and TNFα levels in IMG^N1/T2-dKD^ cells did not differ between stimuli (Figure 6I), indicating that Trem2 is required for cytokine release after changes in microglial cellular state. To extrapolate the impact of NFAT1 on the production of these cytokines, we treated WT^GFP^ and *Neu1*^-/-GFP^ microglia with verapamil. Verapamil decreased the TNFα release by WT^GFP^ and *Neu1*^-/-GFP^ microglia but reduced CCL3 release only by *Neu1*^-/-GFP^ microglia (Figure S6K and S6L). Therefore, *Neu1*^-/-GFP^ microglia are more dependent on NFAT1 signaling than are WT^GFP^ microglia.

Survival of activated microglia relies on Akt/Gsk3β (glycogen synthase kinase 3 beta) antiapoptotic signaling, another pathway targeted by Trem2-FL and sTrem2.^17,29^ Therefore, in stimulated cells, we measured Akt activation as the ratio of phosphorylated Akt at S473 to total Akt (pAkt:totAkt). Under homeostatic conditions, Akt activation was comparable between IMG^NT^ cells and IMG^Neu1-KD^ cells, but under pro-inflammatory conditions, IMG^Neu1-KD^ cells had the highest Akt activation (Figure S6M). IMG^Neu1-KD^ cells maintained higher levels of activated Akt than did IMG^NT^ cells, suggesting that Neu1 downregulation constitutively activates Akt. When stimulation was shifted, Akt activation increased in IMG^NT^ and IMG^Neu1-KD^ cells but decreased in IMG^N1/T2-dKD^ cells (Figure 6J), which demonstrated that Akt activation was Trem2-dependent in these cells. We then measured the inhibition of Gsk3β by calculating the ratio of protein phosphorylated at S9 to total protein (pGsk3β:totGsk3β). IMG^NT^ cells consistently maintained the highest levels of inhibited Gsk3β, (Figure S6N). IMG^Neu1-KD^ and IMG^N1/T2-dKD^ cells showed no difference under homeostatic conditions, but Gsk3β inhibition was higher in IMG^Neu1-KD^ cells under pro-inflammatory stimulation (Figure S6N). Thus, Neu1 appears to modulate Gsk3β inhibition, regardless of microglia cellular state. When stimulation was shifted, Gsk3β inhibition increased only in IMG^Neu1-KD^ cells (Figure 6J). Thus, Neu1 modulates Trem2-dependent Akt/Gsk3β antiapoptotic signaling after microglial activation. Akt activation and subsequent Gsk3β inhibition in IMG^Neu1-KD^ cells was most likely due to sTrem2.

Considering these results, we can infer two concepts: First, in IMG^NT^ cells under pro-inflammatory conditions, Trem2-FL signaling is downregulated, NFκB activation is promoted, and NFAT1 signaling occurs. NFκB and NFAT1 promote the production of TNFα and CCL3. Second, in IMG^Neu1-KD^ cells, Trem2-FL and NFκB signaling are inefficient, but NFAT1 and Akt activation are perpetuated by sTrem2. Akt prevents apoptosis and NFAT1 serves as a compensatory mechanism, promoting the production of TNFα to a lesser extent and that of CCL3 to a greater extent than in IMG^NT^ cells.

### Reinstatement of Neu1 restores homeostatic signaling in deficient microglia

We used our previously described Adeno-associated virus (AAV)-mediated gene therapy to ameliorate the neuroinflammatory phenotype of *Neu1*^-/-^ and AD mouse model hippocampi.^11,34^ We co-injected vectors expressing human Neu1 (scAAV2/8-CMV-huNeu1) and human protective protein/cathepsin A (scAAV2/8-CMV-huPPCA) at a 2:1 ratio into the left-brain hemisphere of mice (Figure 7A and 7B).The expression of the human proteins improved facial morphology, reduced edema, and normalized systemic organ size, which most likely contributed to the increased survival of injected *Neu1*^-/-GFP^ mice (AAV-*Neu1*^-/-GFP^) (Figure S7A-S7C).

**Figure 7.**
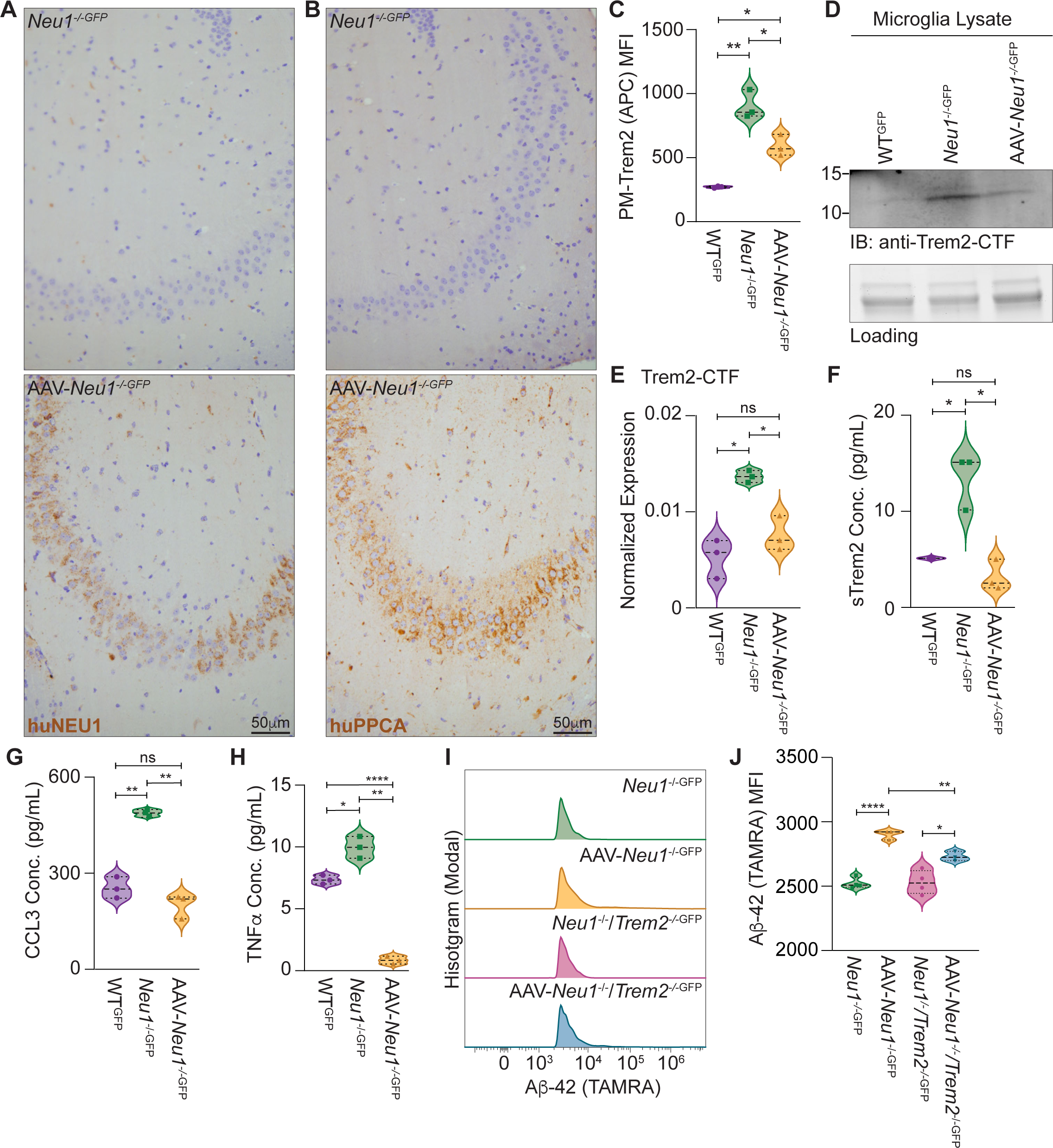
Reinstatement of Neu1 restores homeostatic signaling in deficient microglia. (A-B) Immunohistochemical staining of (A) huNeu1 (brown) and (B) huPPCA (brown) in the CA3 region of 4-month-old *Neu1*^-/-^ and AAV-*Neu1*^-/-^ hippocampi. (C) Quantification of PM-Trem2-FL levels in 4-month-old WT^GFP^, *Neu1*^-/-GFP^, and AAV-*Neu1*^-/-GFP^ microglia by FACS (n=3). (D-E) Representative immunoblot (D) and quantification (E) of Trem2-CTF in 4-month-old WT^GFP^, *Neu1*^-/-GFP^, and AAV-*Neu1*^-/-GFP^ microglia (n=3). (F-H) The concentrations of (F) sTrem2, (G) CCL3, and (H) TNFα in media from WT^GFP^, *Neu1*^-/-GFP^, and AAV-*Neu1*^-/-GFP^ microglia cultures (n=3). (I-J) Representative plots (I) and quantification (J) of TAMRA-labeled Aβ-42 oligomer phagocytosis by *Neu1*^-/-GFP^ (n=3), AAV-*Neu1*^-/-GFP^ (n=3), *Neu1*^-/-^/*Trem2*^-/-GFP^ (n=4), and AAV-*Neu1*^-/-^/*Trem2*^-/-GFP^ (n=3) microglia. Data represented as the medians and quartiles; ns: not significant, *p <0.05, **p <0.01, ****p <0.0001. See also Figure S7.

AAV-*Neu1*^-/-GFP^ microglia were smaller than untreated microglia, suggesting a shift towards ramified morphology (Figure S7D and S7E). PM-Trem2 levels were lower in AAV-*Neu1*^-/-GFP^ microglia than in *Neu1*^-/-GFP^ cells, but they were still higher than that in WT^GFP^ microglia, indicating that Trem2 properties changed after Neu1 re-expression (Figures 7C and S7F). Trem2-CTF levels and sTrem2 release were reduced to WT^GFP^ levels in AAV-*Neu1*^-/-GFP^ microglia (Figure 7D-7F). Thus, Neu1-mediated modulation of Trem2 sialic acid content is required for proper receptor function. AAV-*Neu1*^-/-GFP^ microglia released less CCL3 and TNFα and phagocytosed more Aβ-42 than did *Neu1*^-/-GFP^ controls (Figure 7G-7J). AAV-injected *Neu1*^-/-^/*Trem2*^-/-GFP^ (AAV- *Neu1*^-/-^/*Trem2*^-/-GFP^) microglia also phagocytosed more Aβ-42 than did noninjected *Neu1*^-/-^/*Trem2*^-/-GFP^ cells, but this increase was smaller than that in AAV-*Neu1*^-/-GFP^ cells (Figure 7I and 7J). Thus, Neu1 regulates Trem2-dependent phagocytosis/endocytosis in microglia.

## Discussion

Microglia are central to brain development, homeostasis, and managing pathogenic processes.^13^ How microglia sense and respond to various stimuli hinges on the regulated activity of cell surface receptors and downstream signaling.^13^ Here we uncovered how the sialidase Neu1 controls microglial cellular state by modulating the sialic acid content of Trem2, a substrate of the enzyme. Earlier studies showed that Neu1 deficiency enhances the levels of sialylated APP and the exocytosis of Aβ-42 in sialidosis mice; it also accelerates plaque deposition in an AD mouse model.^11^ These findings suggest a link between Neu1 and AD pathology. We now revealed that the loss of Neu1 activity in microglia, which leads to impaired de-sialylation of Trem2, affected the cycling of the receptor through the endolysosomal system, its proteolytic processing, interactions with DAP12, and signal transduction. These events elicited a toxic, Trem2-dependent, microglia-mediated immune response to AD-related stimuli (e.g., Aβ-42 oligomers).

These findings broaden our understanding of the pathogenic course in a wide range of neurologic conditions that involve lysosomal dysfunction and Trem2-dependent, microglia-mediated neuroinflammation.^19,35–60^ By data mining publicly available GEO (Gene Expression Omnibus) data sets, we identified a significant *Neu1* downregulation in patients with these conditions, including AD, frontotemporal lobar dementia, multiple sclerosis, schizophrenia, bipolar disorder, autism, substance abuse, or normal aging (Table S5). We propose that Neu1 downregulation may contribute to the neuropathogenesis of these diseases/disorders.

### Neu1 is an immunomodulatory molecule in microglia

Neu1 is an immunomodulatory molecule, particularly in cells of the monocytic lineage.^6,7,61–65^ It regulates the differentiation, polarization, and function of various membrane glycoproteins in macrophages; however, our understanding of Neu1 function in microglia remains limited. In microglia cell lines, Neu1 influences TLR4-dependent production of IL-6 and promotes IgG-mediated phagocytosis.^66,67^ In zebrafish, Neu1 deficiency increases *il1β* expression.^68^ The *Neu1*^-/-^ mouse model used in this study enabled us to interrogate the effects of the loss of Neu1 activity in microglia *in vivo*. One key finding is that the production and release of TNFα was promoted by a Neu1-dependent Trem2–NFAT1 signaling mechanism in the absence of NFκB activation. We implicated Neu1 in IL-1α expression and in CCL3 expression, production, and release via lysosomal exocytosis. Because IL-1β is also released via lysosomal exocytosis, that mechanism emerged as a common means by which Neu1 regulates cytokine release from microglia.

Cytokines help regulate astrocyte activation, neuronal survival, and memory formation in the hippocampus.^69–73^ Therefore, alterations in the level of these cytokines, downstream of decreased Neu1 activity, may contribute to cognitive decline, reactive astrocyte activation, and neurotoxicity in normal aging or neurodegenerative diseases. Here we showed that Neu1 controls Aβ-42 oligomer uptake at the level of receptor modification. Thus, Neu1 is a master regulator of microglial phagocytosis. As such, losing Neu1 function could drive neuropathic dysregulation of phagocytosis in various neurodegenerative diseases.

Increased Neu1 levels in macrophages enhance pro-inflammation, and Neu1 downregulation curtails those responses.^74^ However, in hippocampal microglia, we showed that Neu1 deficiency/downregulation perpetuated pro-inflammation rather than dampening it. To fully decipher Neu1’s role as an immunomodulatory molecule and identify potential therapeutic targets, further studies are warranted that will elucidate Neu1-dependent mechanisms regulating cytokine production/release, cellular responses to extracellular cues, and receptor-mediated phagocytosis.

### Trem2 functions and its regulation by Neu1 in microglia

Trem2 became a prominent receptor of interest when it was associated with AD pathology and therapeutics. Early studies on Trem2 revealed the importance of Trem2-FL–DAP12– Syk complexes for initiating phagocytosis, promoting cell survival, and stimulating cytokine production.^19,37,43^ However, few studies have focused on how posttranslational modifications affect Trem2 function. One study has shown that glycosylation of Trem2-FL at Asn20 is required for its trafficking to and signal transduction at the PM, whereas glycosylation at Asn79 is required for signal transduction.^25^ Here we focused on the impact of Trem2-FL sialylation, which we showed does not inhibit its trafficking to the PM but promotes its accumulation at the PM and in endolysosomes. Moreover, sialylation influences the interactions of Trem2 with DAP12 and consequently its ability to induce Syk. Therefore, sialylation of Trem2 acts as a gatekeeper of all Trem2 functions.

In response to Aβ-42 oligomers and other stimuli, Trem2-FL modulates the production of IL-6, IL-1β, and TNFα by regulating their transcription levels.^17,21,23^ We demonstrated that IL-1α expression is also Trem2-dependent, and Neu1 supersedes Trem2 in regulating IL-6 expression and CCL3 and TNFα production. Moreover, Trem2 regulates cytokines at a secondary tier by promoting their release via lysosomal exocytosis in a Neu1-dependent manner. This novel mechanism may represent a therapeutic target, given the potential benefits of modulating lysosomal exocytosis in certain diseases.

Little is known about Trem2-CTF and sTrem2 functions. We have identified Neu1 downregulation as a disease-associated physiological change that promotes Trem2-CTF–mediated inhibition of NFκB. Although sTrem2 promotes Ca^2+^ influx,^30^ this has yet to be connected to sTrem2-dependent signaling. We found that sTrem2 promotes Ca^2+^-dependent NFAT1 activation and is released via Ca^2+^-dependent lysosomal exocytosis. This pathogenic mechanism most likely perpetuates sTrem2 release in diseased brains in which Neu1 is downregulated in microglia. Overall, Trem2 is a prolific receptor with complicated downstream signaling, compounded by the independent functions of its cleaved products Trem2-CTF and sTrem2. Given the impact of Trem2-FL sialylation and the importance of Trem2 function in health and disease states, posttranslational modifications to Trem2-FL, Trem2-CTF, and sTrem2 should be investigated to fully understand how they affect Trem2 signaling.

### Limitations of this study

We limited our studies to the role of Neu1 in hippocampal microglia. However, future studies should investigate whether Neu1 deficiency differentially affects microglia in other brain regions. Due to technical limitations, we could not reliably assess sTrem2 sialylation, which would require advanced glycan analysis methods. Given the complexity of the complement cascade, a comprehensive analysis of the role of Neu1 in this pathway was beyond the scope of this study. However, the robust upregulation of complement gene expression, which parallels findings in other lysosomal storage diseases,^75–79^ motivates further investigation into the role of lysosomes in the classical complement cascade.

## Acknowledgements

A.d’.A. holds the Jewelers for Children Endowed Chair in Genetics and Gene Therapy. This work was funded by NIH grants R01GM104981-04 and 1RF1NS123174-01, The Assisi Foundation of Memphis, and the American Lebanese Syrian Associated Charities (ALSAC). The content is solely the responsibility of the authors and does not necessarily represent the official views of the National Institutes of Health. We thank Drs. Richard Ashmun and Stacie Woolard and the members of the Flow Cytometry and Cell Sorting Core Facility for sorting microglia, which is partially supported by NIH grant P30CA021765; Drs. Aaron Taylor, Aaron Pitre, and Nicolas Denans and the members of the Cell and Tissue Imaging Center–Light Microscopy Shared Resource for their assistance with processing and imaging tissue and microglia; and Dr. Geoffrey Neale and the members of the Hartwell Center for the microarray analyses. We also thank Drs. Angela McArthur and Gerard Grosveld for their review and editing of the finalized manuscript.

## Author Contributions

Conceptualization, L.E.F. and A.d’A. Methodology, L.E.F., H.H., I.A., and A.d’A. Validation, L.E.F. and A.d’A. Formal Analysis, L.E.F. and J.A.W. Investigation, L.E.F., I.A., and A.d’A. Resources, L.E.F., D.v.d.V., H.H., E.G., and A.d’A. Data Curation, L.E.F. and A.d’A. Writing – Original Draft Preparation, L.E.F. Writing – Review & Editing, L.E.F. and A.d’A. Visualization, L.E.F., D.v.d.V., and A.d’A. Supervision, A.d’A. Project Administration, A.d’A. Funding Acquisition, A.d’A. All authors have read and agreed to the published version of the manuscript.

## Declaration of interest

The authors declare no conflict of interest.

## STAR Methods

### Key Resources Table

### Resource Availability

#### Lead contact

Further information and requests for resources and reagents should be directed to and will be fulfilled by the lead contact, Alessandra d’Azzo.

#### Materials availability

Lentiviruses encoding the shRNAs used in this study are readily available at Horizon Discovery Dharmacon™. This study did not generate any unique new reagents.

#### Data and code availability

- Microarray data have been deposited to the NCBI GEO databank.
- GSEA data have been deposited to the NCBI GEO databank.
- No original code was generated for this study.
- Any additional information required to re-analyze the data reported here is available from the Lead Contact upon request.

### Experimental Models Details

#### Animal Models

Animals were housed in a fully AAALAC (Assessment and Accreditation of Laboratory Animal Care)-accredited animal facility with controlled temperature (22°C), humidity, and lighting (alternating 12-h light/dark cycles). Food and water were provided *ad libitum*. All procedures in mice were performed according to animal protocols approved by the St. Jude Children’s Research Hospital Institutional Animal Care and Use Committee and NIH guidelines. Sexes were represented equally. WT and *Neu1^-/-^* mice were bred into the C57BL/6J background. CX3CR1^GFP^ mice were obtained from Jackson Laboratories and crossed with WT and *Neu1^-/-^* animals in the C57BL/6J background to produce WT^GFP^ and *Neu1^-/-^*^GFP^ mice. *Trem2^-/-^* mice were obtained from Jackson Laboratories and bred into the C57BL/6J background. *Neu1^-/-^*/*Trem2^-/-^*^GFP^ and *Trem2^-/-^*^GFP^ mice were generated by crossing *Neu1^+/-^*^GFP^ and *Trem2^-/-^* mice, which were obtained from Jackson Laboratories, in the C57BL/6J background.

#### Cell lines

Immortalized microglia, derived from 8-week-old female C57Bl6 mice infected with v-raf/v-myc retrovirus, were purchased from Sigma and transduced at a multiplicity of infection of 10 with lentiviral vectors encoding RFP-labeled nontargeting shRNA, RFP-Neu1–targeting shRNA, or RFP-Neu1– and GFP-Trem2–targeting shRNAs to produce IMG^NT^, IMG^Neu1-KD^, and IMG^N1/T2-dKD^ cells, respectively, in DMEM containing 10% fetal bovine serum (FBS), PenStrep, GlutaMAX, and 8 μg/mL polybrene for 24 h at 37°C in 5% CO_2_. To establish stable cell lines, newly transduced cells were grown in DMEM containing 10% FBS, PenStrep, GlutaMAX, and 10 μg/mL puromycin at 37°C in 5% CO_2_ until confluent in a T150 flask. The cells were then washed, suspended in FACS buffer, and sorted by FACS for the top 10% highest fluorescing population (RFP for IMG^NT^ and IMG^Neu1-KD^ cells; RFP/GFP for IMG^N1/T2-dKD^ cells), which was subsequently grown to confluency in a 10-mm plate in DMEM containing 10% FBS, PenStrep, and GlutaMAX at 37°C in 5% CO_2_. Cells in complete DMEM with 10% DMSO were frozen and stored in liquid nitrogen until needed.

#### Primary Cultures

Primary microglia were isolated from female and male WT*^GFP^*, *Neu1^-/-GFP^*, *Neu1^-/-^* /*Trem2^-/-GFP^*, and *Trem2^-/-GFP^*hippocampi and were maintained in culture in Microglia Media at 37°C in 5% CO_2_.

### Methods Details

#### Immunofluorescence Analyses

For hippocampal tissue analyses, mouse brains were collected and stored in 10% formalin until embedding in paraffin. Immunofluorescence (IF) analyses were performed on 6-μm-thick serial paraffin sections. Sections were subjected to automated de-paraffinization and antigen retrieval using a 20-min cycle in a rice cooker with slides submerged in 10 mM sodium citrate/0.1% Tween-20. After permeabilization for 45 min in 0.5% TritonX-100/PBS and blocking for 1 h in 2% BSA/0.5% TritonX-100/PBS, sections were incubated overnight at 4°C with anti-Iba1 antibody diluted (1:1000) in blocking buffer. Sections were then washed 3 times for 5 min in blocking buffer and subsequently incubated with the anti-rabbit IgG-AF488 diluted (1:400) in blocking buffer for 2 h at room temperature (RT). Sections were washed twice for 5 min in blocking buffer, incubated in a solution containing DAPI diluted (1:1000) in PBS for 10 min at RT, washed for 5 min in PBS, and mounted using ProLong*^TM^*Gold antifade. Sections were imaged using an AxioScan microscope in the Light Microscopy Core at St. Jude Children’s Research Hospital. Images were analyzed using Zen software.

For microglia culture analyses, microglia were isolated from WT*^GFP^*, *Neu1^-/-GFP^*, *Neu1^-/-^/Trem2^-/-GFP^*, and *Trem2^-/-GFP^* brains, plated at 150,000 cells/well in 24-well plates in 1.5 mL microglia media, and left to adhere for 4 h at 37°C, 5% CO_2_. Following adhesion, media was replaced with 1 mL fresh Microglia Media and left to culture at 37°C, 5% CO_2_ overnight. Cells were subsequently fixed in 4% PFA/PHEM, permeabilized with 0.5% Tween-20/PHEM, and subjected to blocking for 1 h at RT by using 0.2% Tween-20/10% serum/PHEM.

After blocking, cells were incubated with gentle rocking at 4°C in 10% serum/PHEM containing SNA-Cy5 and antibodies against Lamp1, CD68, Golgin97, EEA1, and Trem2. Cells were imaged as Z-stacks at 3× zoom using a 63× oil lens on a Leica SP8 microscope. Images were converted into Imaris files and analyzed for cell volume and spot volume of the various markers by using the Imaris cell and spot functions. Co-labeling of markers was quantified by object-intensity co-localization using the Imaris spot function.

#### Immunohistochemical Analyses

Mouse brains were collected and kept in 10% formalin until paraffin embedding. Immunohistochemical (IHC) analyses were performed on 6-μm-thick serial paraffin sections. Sections were subjected to automated de-paraffinization and antigen retrieval using a 20-min cycle in a pressure cooker with slides submerged in 10 mM sodium citrate/0.1% Tween-20. After washing sections 3 times for 5 min in PBS, they were blocked in 0.5% Tween-20/10% normal serum/PBS for 2 h at RT then incubated overnight at RT with antibodies against Iba1 (1:1000), human Neu1 (1:100), or human PPCA (1:200) diluted in blocking buffer. The sections were then washed 3 times for 5 min with PBS and incubated with biotinylated secondary antibody for 2 h at RT. Endogenous peroxidase was quenched by incubating the sections with 5% hydrogen peroxidase/PBS for 15 min at RT. Antibodies were detected using the VECTASTAIN^®^ Elite^®^ ABC Kit, and diaminobenzidine substrate, and sections were counterstained with hematoxylin per standard methods.

#### Flow Cytometry Analyses

Brain mononuclear cells (BMCs) were isolated from WT^GFP^, *Neu1^-/-^*^GFP^, *Trem2^-/-^*^GFP^, *Neu1^-/-^*/*Trem2^-/-^*^GFP^, and AAV-*Neu1^-/-^*^GFP^ hippocampi and stained with anti–Trem2-APC and anti–Lamp1-PE in FACS buffer (10% FBS/2 mM EDTA/PBS) for 20 min at RT in the dark. Cells were washed, re-suspended in FACS buffer containing 1:10 DAPI, and analyzed using an LSRFortessa^TM^ flow cytometer. Data were analyzed and mean fluorescence intensity (MFI) was calculated using FlowJo software.

#### Microarray Analyses

Total RNA (100 ng) was extracted from WT and *Neu1^-/-^* hippocampi at 1 and 5 months, converted into biotin-labeled cRNA, and hybridized to a mouse GeneChip. Probe signals from microarrays were normalized and transformed into log_2_ transcript expression values by using the Robust Multiarray Average algorithm. Differentially expressed transcripts were identified by ANOVA, and the false discovery rate (FDR) was estimated. GSEA were completed using xtools software.

#### Quantitative Real-time PCR Analyses

Total RNA was isolated from 2- and 4-month-old WT and *Neu1^-/-^* hippocampi or 4-month-old WT^GFP^, *Neu1^-/-^*^GFP^, *Neu1^-/-^*/*Trem2^-/-^*^GFP^, and *Trem2^-/-^*^GFP^ hippocampi by using the PureLink RNA kit per the manufacturer’s protocol. DNA contaminants were removed using a DNAse I column per the manufacturer’s protocol. RNA quantity and purity were measured using a Nanodrop Lite spectrophotometer. Complementary DNA was produced using 24 μg total RNA with RT_2_ First Strand Kit. RT-qPCR was performed using the RT_2_ Sybr Green qPCR Mastermix, 1-5 μL (50-250 ng) cDNA, 10 μM primer, and RNAse-free water in a 25-μL reaction volume on a CFX96 real-time PCR machine. Samples were normalized using 18S ribosomal RNA expression.

#### β-Hexosaminidase Activity Assays

Media from 4-month-old WT, *Neu1^-/-^* WT^GFP^, *Neu1^-/-^*^GFP^, *Neu1^-/-^*/*Trem2^-/-^*^GFP^, and *Trem2^-/-^*^GFP^ microglia in culture were collected and centrifuged at 11,000 ×*g* for 5 min. Spun-down medium was applied onto a G50 Sephadex column packed in an acidic buffer before performing enzymatic assays of β-hex activity. Medium (10 μL) was incubated with 10 μL 4MU-N-acetyl-β-D-glucosaminide. To stop enzyme reactions, 200 μL 0.5 M carbonate buffer (pH 10.7) was added to all wells. Fluorescence was measured on a plate reader (EX-355 nm, EM-460 nm). The net fluorescence values were compared with those of the linear 4MU standard curve and were used to calculate the specific enzyme activities. Activities were calculated as nanomoles of substrate converted per hour per milligram of protein.

#### CSF collection

For collection of CSF, 4-month-old WT^GFP^ and *Neu1^-/-^*^GFP^ mice were anaesthetized with avertin and placed into a Kopf stereotaxic device. A small incision was made at the base of the skull, and CSF was collected from the cisterna magna by using a syringe attached to a butterfly needle. Collected CSF was dispensed into 250 μL Eppendorf tubes, frozen in liquid nitrogen, and then stored at –80°C until needed.

#### Cycloheximide Treatments and CSF Stimulation of IMG Cell Cultures

Established IMG^NT^, IMG^Neu1-KD^, and IMG^N1/T2-dKD^ cell lines were plated at 2×10^6^ cells/plate in 10-cm plates in DMEM containing 10% FBS, PenStrep, GlutaMAX, for 24 h at 37°C in 5% CO_2_. Plates were washed with PBS, and media was replaced with Microglia Media for 24 h at 37°C in 5% CO_2_. Plates were then washed with PBS, and media was replaced with treatment media, as described below.

For CHX assays, media was replaced with Microglia Media containing 10 μM CHX, alone or in combination with 20 μM GM6001, a ubiquitous inhibitor of metalloproteinases, or 25 nM BafA1, a potent inhibitor of lysosomal exocytosis, and incubated at 37°C in 5% CO_2_ for 0.25, 0.5, 1, 2, or 4 h; untreated cells were used as 0 h controls. At the appropriate time point, cells were washed with PBS and lysed in the culture dish in CST lysis buffer containing protease, phosphatase, and GM6001 inhibitors. Protein concentrations were determined using bicinchoninic acid (BCA) assay, then samples containing 30 μg total protein were prepared for immunoblot analysis.

For CSF stimulation assays, media was replaced with Microglia Media containing CSF collected from 4-month-old WT or Neu1 microglia (diluted 1:500) and incubated at 37°C in 5% CO_2_ for 24 h. Cells were then washed with PBS and lysed in the culture dish in CST lysis buffer containing protease, phosphatase, and GM6001 inhibitors. Protein concentrations were determined using BCA assay, and then samples containing 30 μg total protein were prepared for immunoblot analysis.

#### Enzyme-linked Immunosorbent Assays

CCL3, TNFα, and sTrem2 were measured in the soluble fraction of 4-month-old WT^GFP^ and *Neu1^-/-^*^GFP^ hippocampal lysates, CSF collected from 4-month-old WT^GFP^ and *Neu1^-/-^*^GFP^ mice, in the supernatant collected from 4-month-old WT^GFP^, *Neu1^-/-^*^GFP^, *Trem2^-/-^*^GFP^, *Neu1^-/-^*/*Trem2^-/-^*^GFP^, and AAV-*Neu1^-/-^*^GFP^ microglia cultures, and media collected from IMG^NT^, IMG^Neu1-KD^, and IMG^N1/T2-dKD^ cultures. The hippocampal soluble fractions were obtained after homogenization of hippocampal tissue. Freshly collected tissue was homogenized in 150 μL cell lysis buffer 2 containing 2× protease inhibitors using a TissueLyzer II set to run at 20 Hz/s for 4 min. Homogenates were then spun at 10,000 ×*g* for 5 min. Supernatants from spun samples were collected into new tubes and labeled as soluble fractions.

After the protein concentration was quantified using a BCA assay, freshly prepared soluble fractions were aliquoted into 150-μg volumes, which were flash-frozen on dry ice and stored at –80°C. To obtain culture supernatants, GFP^hi^ microglia were isolated from total brains and plated at 150,000 cells/well in a 24-well plate in 1.5 mL Microglia Media. Cells were allowed to attach for 4 h at 37°C and 5% CO_2_. Following cell attachment, the media was removed and replaced with 1 mL fresh Microglia Media. Cells were maintained in culture for 24 h at 37°C, 5% CO_2_, at which point supernatants were collected in 50-μL aliquots, frozen on dry ice, and stored at –80°C for future use. Alternatively, cells were maintained in culture for 20 h at 37°C, 5% CO_2_, at which point vacuolin-1, an inhibitor of lysosomal exocytosis, was added to a final concentration of 10 μM, or verapamil, a Ca^2+^ chelator that inhibits NFAT1, was added to a final concentration of 50 μM, and cells were maintained in culture for an additional 4 h.

CCL3 and TNFα levels in tested samples were determined using the sandwich enzyme-linked immunosorbent assay (ELISA) kits per the manufacturer’s instructions. Briefly, 100 μL/well of 1:1 assay diluent:test sample or assay diluent:standard was added to 96-well plates and incubated for 2 h. Plates were washed 5 times with 400 μL wash buffer and then incubated for 2 h with 100 μL/well of the conjugate solution. Plates were rinsed 5 times with wash buffer and incubated in the dark for 30 min with 100 μL/well of substrate solution. After this period, 100 μL/well of stop solution was added to terminate the reaction. The absorbance was measured at 450 nm, with a correction absorbance measured at 540 nm. For sTrem2 ELISAs, 100 µL standard or sample (diluted accordingly in assay diluent B) was added to each well and incubated for 2.5 h at RT. Following this, plates were washed 4 times, and then 100 µL prepared biotin antibody was added to each well and incubated for 1 h at RT. Next, plates were washed 4 times and 100 µL prepared streptavidin solution was added to each well and incubated for 45 min at RT. Then, plates were washed 4 times and 100 µL TMB One-Step Substrate Reagent was added to each well and incubated for 30 min at RT in the dark. Finally, without washing, 50 µL stop solution was added to each well. Plates were read at 450 nm immediately.

#### *In Vivo* Phagocytosis Assays

Phagocytosis materials were prepared as follows: 1.0 μm fluorescent red, amine-modified, latex beads were diluted 1:5 in FBS, incubated at 37°C for 1 h, and then diluted (1:10) in sterile PBS. AF594-Zymosan A BioParticles^TM^ were diluted in 2 mM NAN_3_ to 5 μg/μL; 200 μM TAMRA-Aβ-42 was diluted (1:20) in sterile PBS to a final concentration of 10 μM and oligomerized at 4°C for 24 h. After preparation of the materials, animals were anaesthetized with Avertin and placed into a Kopf stereotaxic device. Injections were done unilaterally into the left hippocampus by using a 10-μL Hamilton syringe fitted with a glass micropipette. A 5-μL sample of the aforementioned material was injected. For all brain injections, the needle was left in place for 5 min before withdrawal. Mice were housed in individual units after the injection and left to recover overnight. Then mice were anesthetized, perfused with ice-cold PBS, and brains were collected for isolation processing. WT^GFP^, *Neu1^-/-^*^GFP^, *Trem2^-/-^*^GFP^, *Neu1^-/-^*/*Trem2^-/-^*^GFP^, and AAV-*Neu1^-/-^*^GFP^ BMCs were isolated, and GFP^hi^ microglia populations were identified and analyzed using a LSRFortessa^TM^ flow cytometer. Data were analyzed and the MFI was calculated using FlowJo software.

#### Immunoblotting

Hippocampi from WT and *Neu1^-/-^* mice, microglia isolated from WT^GFP^, *Neu1^-/-^*^GFP^, *Trem2^-/-^*^GFP^, and AAV-*Neu1^-/-^*^GFP^ mice, and IMG^NT^, IMG^Neu1-KD^, and IMG^N1/T2-dKD^ lysates were homogenized with CST lysis buffer or CHAPS buffer containing protease, phosphatase, and GM6001 inhibitors. Protein concentrations were determined using a BCA assay. Proteins were separated by SDS–PAGE on precast Mini-PROTEAN 4%-20%, or Criterion Midi 4%-20%, 10%, or 12% TGX gels under reducing conditions and transferred to a polyvinylidene difluoride membrane.

Membranes were incubated for 1 h in blocking buffer (5% dry milk in TBS-Tween) at RT and subsequently probed with antibodies against Trem2 (1:500), C-terminal Trem2 (1:1000), DAP12 (1:1000), Syk (1:1000), pSyk^Y525/526^ (1:1000), NFκB (1:1000), pNFκB^S536^ (1:1000), calcineurin (1:1000), NFAT1 (1:1000), Akt (1:1000), pAkt^S473^ (1:1000), Gsk3β (1:1000), and pGsk3β^S9^ (1:1000) diluted in 5% BSA/TBS-Tween overnight at 4°C. Following this step, blots were washed 3 times for 5 min with TBS-Tween and then incubated with the appropriate secondary antibody for 2 h at RT. Immunoblots were developed using the Clarity Max^TM^ Western ECL substrate.

#### Isolation of Brain Mononuclear Cells

Hippocampi from 2-month-old WT^GFP^ and *Neu1^-/-^*^GFP^ mice or from 4-month-old WT^GFP^, *Neu1^-/-^*^GFP^, *Neu1^-/-^*/*Trem2^-/-^*^GFP^, *Trem2^-/-^*^GFP^, and AAV-*Neu1^-/-^*^GFP^ mice were collected into DMEM on ice. Tissue was added to 2-5 mL digestion buffer (papain/EDTA/0.01% collagenase type II/50 μg/mL DNase in RPMI) in gentleMACS C-tubes then run on a MACS Dissociator for 30 min at 37°C, 28 rpm. Digested tissues were spun briefly at 300 ×*g*, filtered through a 70-μm nylon cell strainer, washed with 10 mL FACS buffer, and then spun at 380 ×*g* for 10 min at 18°C. Pellets were resuspended in 6-8 mL 30% Percoll^®^ and spun down for 30 min at 300 ×*g* at 18°C. The top 2 layers (myelin and supernatant) were removed, and the BMCs were washed in 6 mL FACS buffer and centrifuged at 450 ×*g* for 7 min at 18°C. The pellets were then resuspended in FACS buffer containing 1:10 DAPI for sorting of GFP^hi^ microglia, phagocytosis assay analyses, or staining for flow cytometry analyses.

#### Plasma Membrane and Crude Lysosomal Fraction Isolation

The PM fractions of 4-month-old WT and *Neu1^-/-^* hippocampi were isolated as previously described.^80^ Mice were euthanized by CO_2_; hippocampi were isolated and homogenized in a 2-mL Kontes Dounce Homogenizer (Kimble) in 1 mL homogenizer solution. Homogenates were diluted 10% w/v with homogenizer solution and centrifuged at 1400 ×*g* for 10 min at 4°C. Pellets were resuspended in 10% w/v homogenizer solution, homogenized, and centrifuged at 710 ×*g* for 10 min at 4°C. Supernatants were isolated in a Sorvall centrifuge at 13,800 ×*g* for 10 min at 4°C. Pellets were resuspended in 10% w/v homogenizer solution, homogenized, and centrifuged at 13,800 ×*g* for 10 min at 4°C. This was repeated twice. Pellets were homogenized in 2 mL 11% w/v sucrose in ER-PM buffer, then homogenates were layered on a discontinuous sucrose gradient and centrifuged at 100,000 ×*g* for 2.5 h at 4°C. The PM band was collected, resuspended in 15 mL ER-PM wash buffer, and centrifuged at 13,000 ×*g* for 10 min at 4°C. Supernatants were centrifuged at 48,000 ×g for 20 min at 4°C. Purified PM fractions were resuspended in CST lysis buffer. Protein concentrations were determined using a BCA assay, and then samples containing 25 μg total protein were prepared for immunoblot analysis.

Lysosome isolation kits were used to isolate the crude lysosomal fraction from 4-month-old WT and *Neu1^-/-^*hippocampi. Mice were euthanized by CO_2_, and hippocampi were isolated and homogenized in 4 volumes 1× extraction buffer and then centrifuged at 1000 ×*g* for 10 min at 4°C. Pellets were resuspended in 2 volumes 1× extraction buffer, homogenized, and centrifuged at 1000 ×*g* for 10 min at 4°C. Supernatants were centrifuged at 20,000 ×*g* for 20 min at 4°C. Crude lysosomal fractions were resuspended in CST lysis buffer. Protein concentrations were determined using BCA assay, and then samples containing 25 μg total protein were prepared for immunoblot analysis.

#### Co-Immunoprecipitation

CSF-stimulated IMG^NT^, IMG^Neu1-KD^, and IMG^N1/T2-dKD^ lysates were homogenized with CHAPS buffer containing protease, phosphatase, and GM6001 inhibitors. Protein concentrations were determined using a BCA assay. For DAP12 immunoprecipitations, 100 μg total lysate was used for each immunoprecipitation. Samples were rotated at 4°C for 16 h with 2 μg anti-DAP12 antibody or 2 μg anti–rabbit IgG isotype control in a final volume of 200 μL CHAPS. After incubation, 40 μL washed protein A magnetic beads diluted (1:2) in CHAPS was added to each sample. Samples were rotated for 1 h at RT and then placed on a magnet. Beads were washed 4 times in CHAPS and then resuspended in 20 μL LDS running buffer and boiled for 5 min at 95°C. Samples where then spun down, placed on a magnet, and loaded on SDS-PAGE gels to be analyzed by immunoblot analysis; samples contained 10 μg total protein for input controls.

For each SNA immunoprecipitation, 50 μg total lysate was used. Samples were rotated at 4°C for 16 h with 2 μg of biotinylated SNA or 2 μL CHAPS buffer for beads only controls, in a final volume of 200 μL CHAPS. After incubation, 10 μL washed Nanolink™ streptavidin magnetic beads, diluted (1:2) in CHAPS, was added to each sample. Samples were rotated for 1 h at RT and then placed on a magnet. Beads were washed 4 times in CHAPS and then resuspended in 20 μL LDS running buffer and boiled for 5 min at 95°C. Samples where then spun down, placed on a magnet, and loaded on SDS-PAGE gels to be analyzed by immunoblot; samples were prepared accordingly, containing 5 μg total protein for input controls.

#### Vector Production

The *scAAV2/8-CMV*-*CTSA* (huPPCA) and *scAAV2/8-CMV*-*Neu1* (huNeu1) constructs contained a cytomegalovirus (CMV) promoter, which ensured the expression of the 1.44-kb human *PPCA* complementary DNA and the 1.247-kb human *Neu1* cDNA. The scAAV vector particles were made in the Children’s GMP, LLC facility on the St. Jude campus. The genome titer of each vector was determined by qPCR and/or direct loading and electrophoresis of detergent-treated vector particles on native agarose gels, staining with fluorescent dye, and quantification of signal relative to known mass standards.

#### AAV Transduction *In Vivo*

Animals were anaesthetized and placed into a Kopf stereotaxic device. Intracerebroventricular injections were done unilaterally into the left hippocampus. Injections were performed with a 10-μL Hamilton syringe fitted with a glass micropipette. 10.25 μL 2 Neu1 GC: 1 PPCA GC virus was injected per mouse (10 μL PPCA virus, titer: 1 × 10^12^ GC and 0.25 μL Neu1 virus, titer: 7.97 × 10^13^ GC). After each injection, the needle was left in place for 5 min before withdrawal from the brain. Mice were housed individually for 3 months after the injection, then anesthetized, perfused with ice-cold PBS, and their brains were processed for downstream analyses.

#### Survival Analysis

A cohort of WT^GFP^ (n=5), *Neu1^-/-^*^GFP^ (n=5), and AAV-*Neu1^-/-^*^GFP^ (n=4) mice were kept past the 4-month experimental end point, until reaching a humane end point or 12 months of age, whichever occurred first. Graphpad Prism 10 software was used to calculate survival using Kaplan-Meier survival analyses. The lifespan of each mouse, in days, was plotted, and survival was presented as the percentage of the total group remaining.

#### Imaging of Mice and Systemic Organs

WT^GFP^, *Neu1^-/-^*^GFP^, and AAV-*Neu1^-/-^*^GFP^ mice were assessed for various physical attributes. Before tissue collection, pictures of the head (side-view) and paws (palm up) were acquired using a Samsung Galaxy S23 camera. The heart, spleen, and kidney of each mouse were isolated and measured with a metric ruler. A Samsung Galaxy S23 camera was used to photograph the organs.

#### Quantification and Statistical Analyses

Statistical analyses were performed using GraphPad Prism 10 software. Quantitative data are represented as mean ± SD or as medians + quartiles. For comparisons between 2 groups, Welch’s student’s *t*-tests (unpaired, 2-tailed) were performed. For comparisons of 3 or more groups, 2-way ANOVA with multiple comparisons was performed. P-values <0.5 were considered statistically significant. The number of replicates for a given experiment is indicated in each figure legend.

#### Key Resources Table

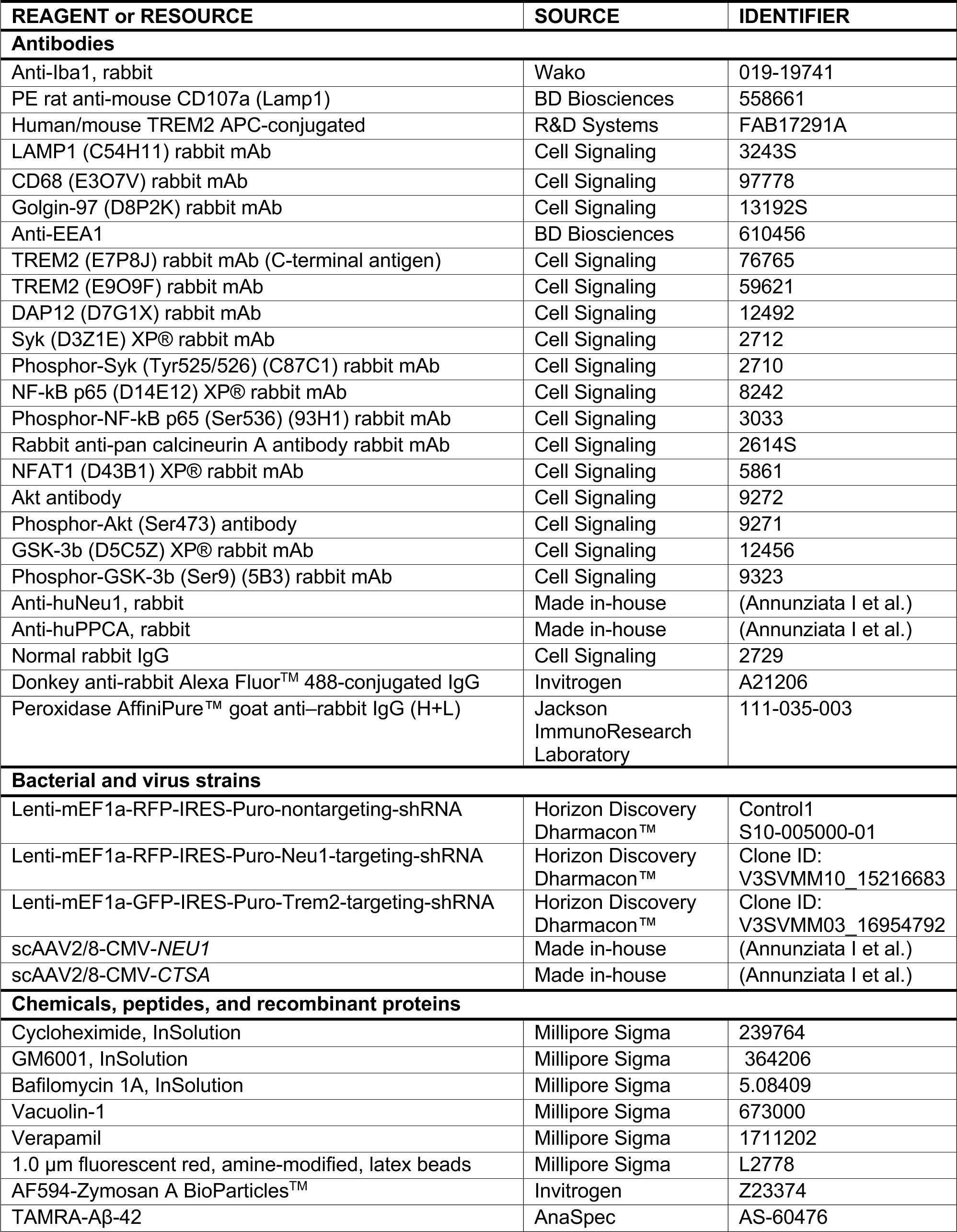
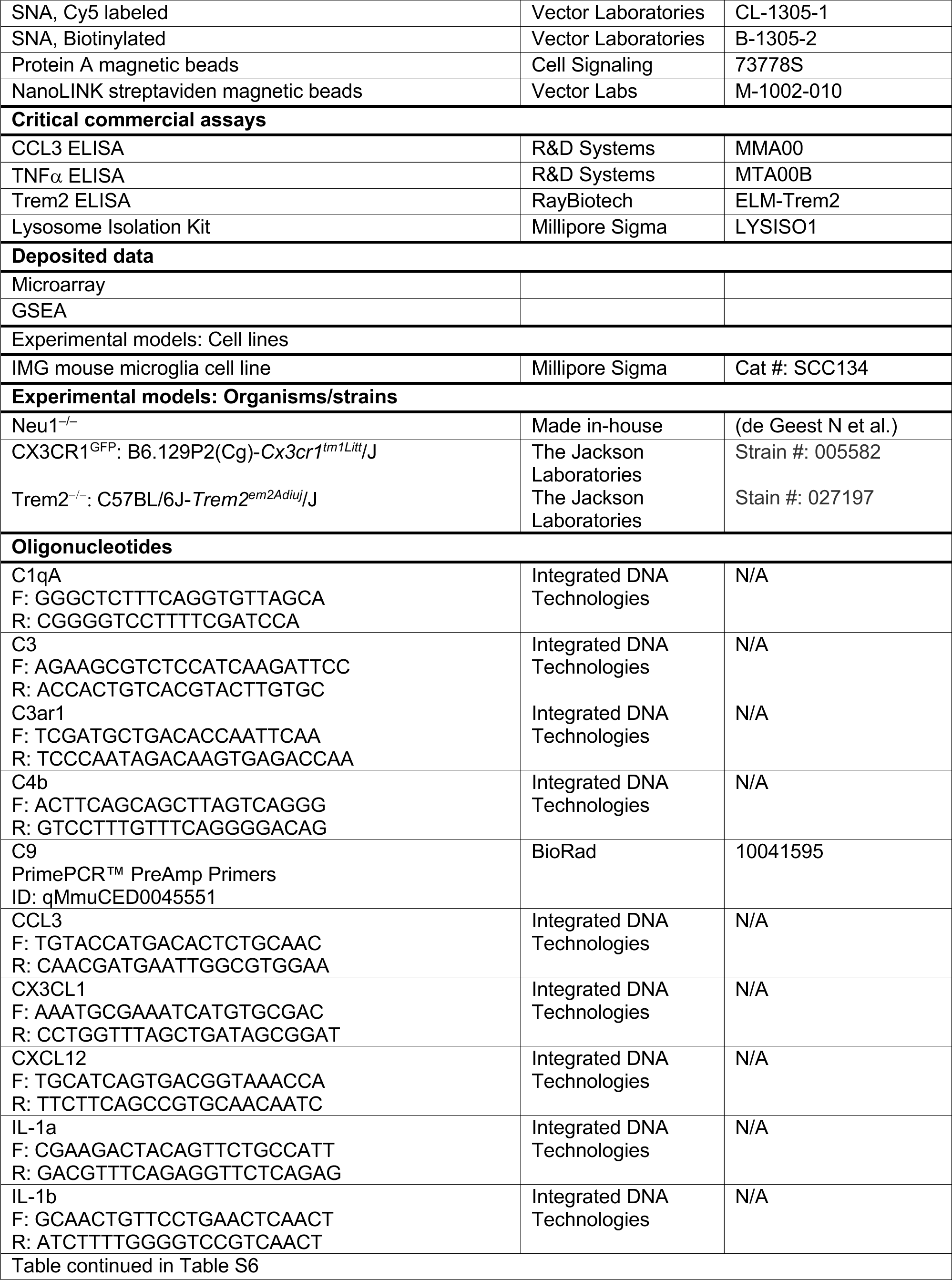
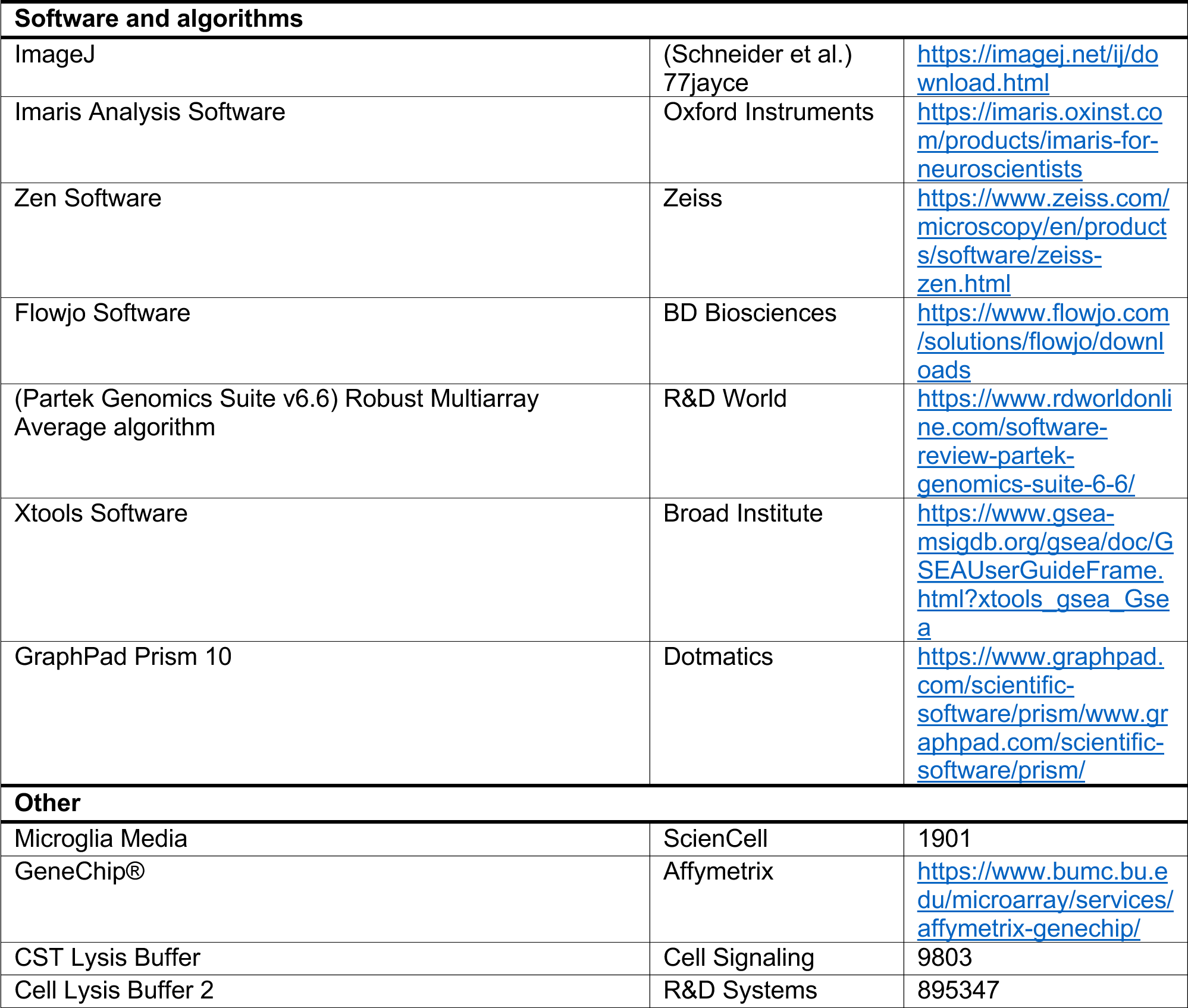

---

**Figure S1.**
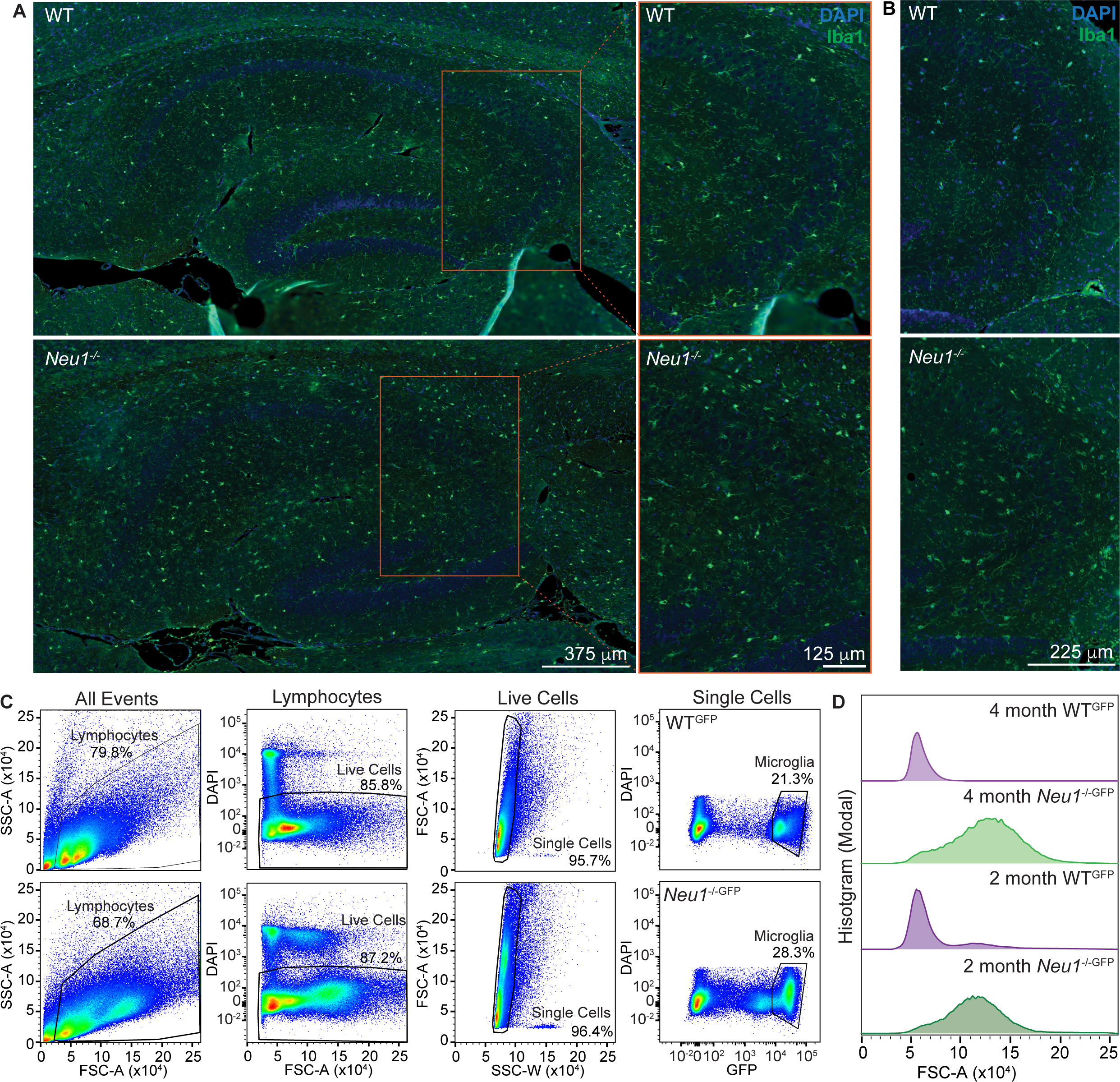
Neu1*^-/-^* microglia have altered morphology. (A) IF staining of microglia in 2-month-old WT and Neu1*^-/-^* hippocampi; orange boxed area depicts staining in the CA3 region; Iba1 (green) and DAPI (blue). (B) IF staining of microglia in the CA3 region of 5-month-old WT and Neu1*^-/-^* hippocampi; Iba1 (green) and DAPI (blue). (C) The gating strategy used to isolate GFP^hi^ microglia for flow cytometry analyses. Representative plots are shown for WT^GFP^ (top) and Neu1*^-/-^*^GFP^ (bottom) samples. (D) Representative plots of cell size (FSC-A) of 2- and 4-month-old WT^GFP^ and Neu1*^-/-^*^GFP^microglia. Related to Figure 1.

**Figure S2.**
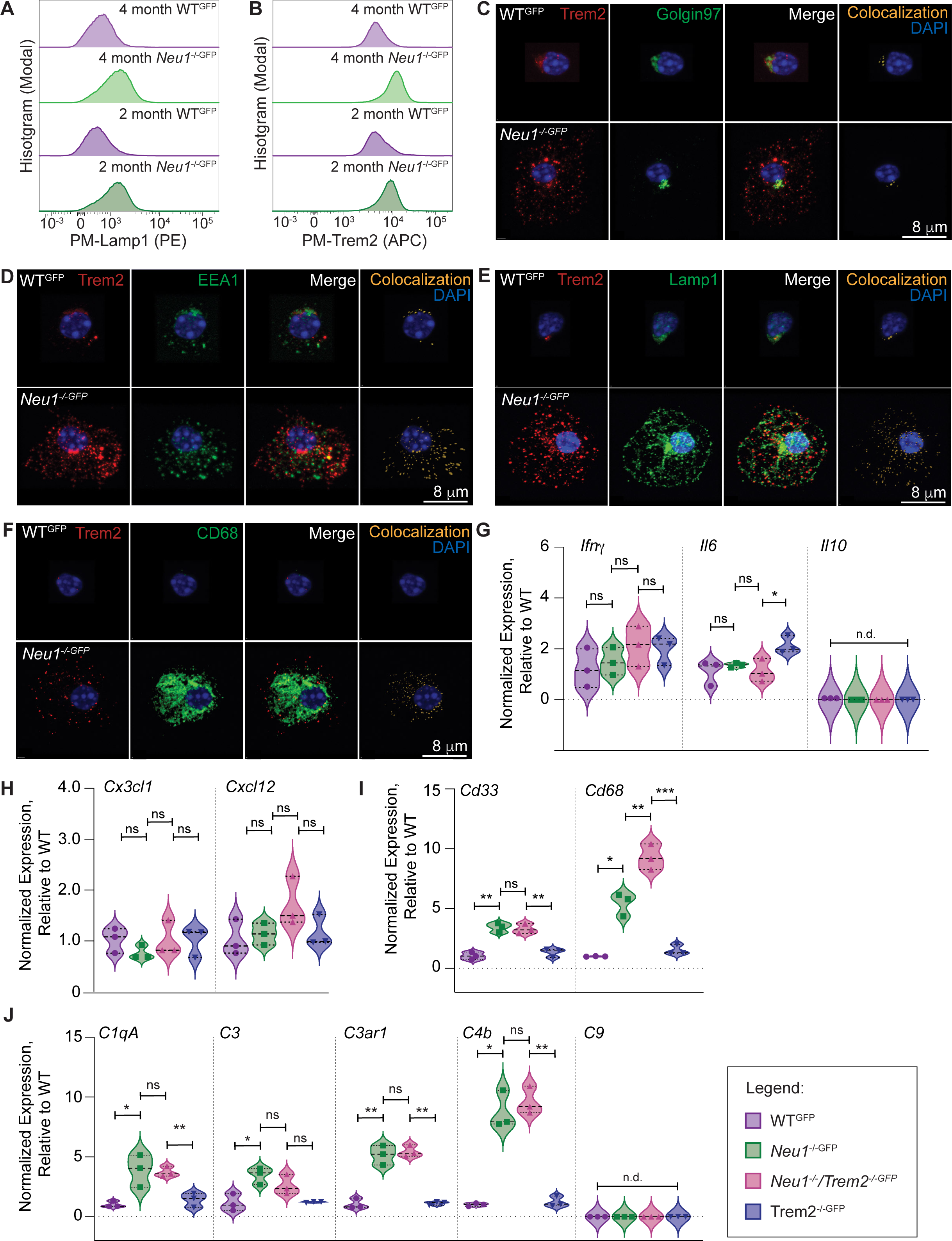
Subcellular localization of Trem2 and genetic profiling of Neu1*^-/-^*/Trem2*^-/-^* hippocampi. (A-B) Representative plots of (A) PM-Lamp1 and (B) PM-Trem2-FL expressed in 2- and 4-month-old WT^GFP^ and Neu1*^-/-^*^GFP^ microglia. (C) Representative images of IF staining depicting golgi-localized Trem2 in 4-month-old WT^GFP^ and Neu1*^-/-^*^GFP^ microglia; Trem2 (red), Golgin97 (green), DAPI (blue), and co-localized objects (yellow) are shown. (D) Representative images of IF staining depicting early endosome-localized Trem2 in 4-month-old WT^GFP^ and Neu1*^-/-^*^GFP^ microglia; Trem2 (red), EEA1 (green), DAPI (blue), and co-localized objects (yellow) are shown. (E) Representative images of IF staining depicting lysosome-localized Trem2 in 4-month-old WT^GFP^ and Neu1*^-/-^*^GFP^ microglia; Trem2 (red), Lamp1 (green), DAPI (blue), and co-localized objects (yellow) are shown. (F) Representative images of IF staining depicting phagosome-localized Trem2 in 4-month-old WT^GFP^ and Neu1*^-/-^*^GFP^ microglia; Trem2 (red), CD68 (green), DAPI (blue), and co-localized objects (yellow) are shown. (G-J) The qRT-PCR analyses of gene expression in 4-month-old WT^GFP^, Neu1*^-/-^*^GFP^, Neu1*^-/-^*/Trem2*^-/-^*^GFP^, and Trem2*^-/-^*^GFP^ hippocampi for (G) Ifn*g*, Il6, and Il10; (H) Cx3cl1, and Cxcl12; (I) Cd33 and Cd68; and (J) C1qa, C3, C3ar1, C4b, and C9 (n=3 for each group). Data represented as the medians and quartiles; n.d., not detected; ns, not significant; *p≤0.05, **p≤0.01, ***p≤0.001. Related to Figures 3 and 4.

**Figure S3.**
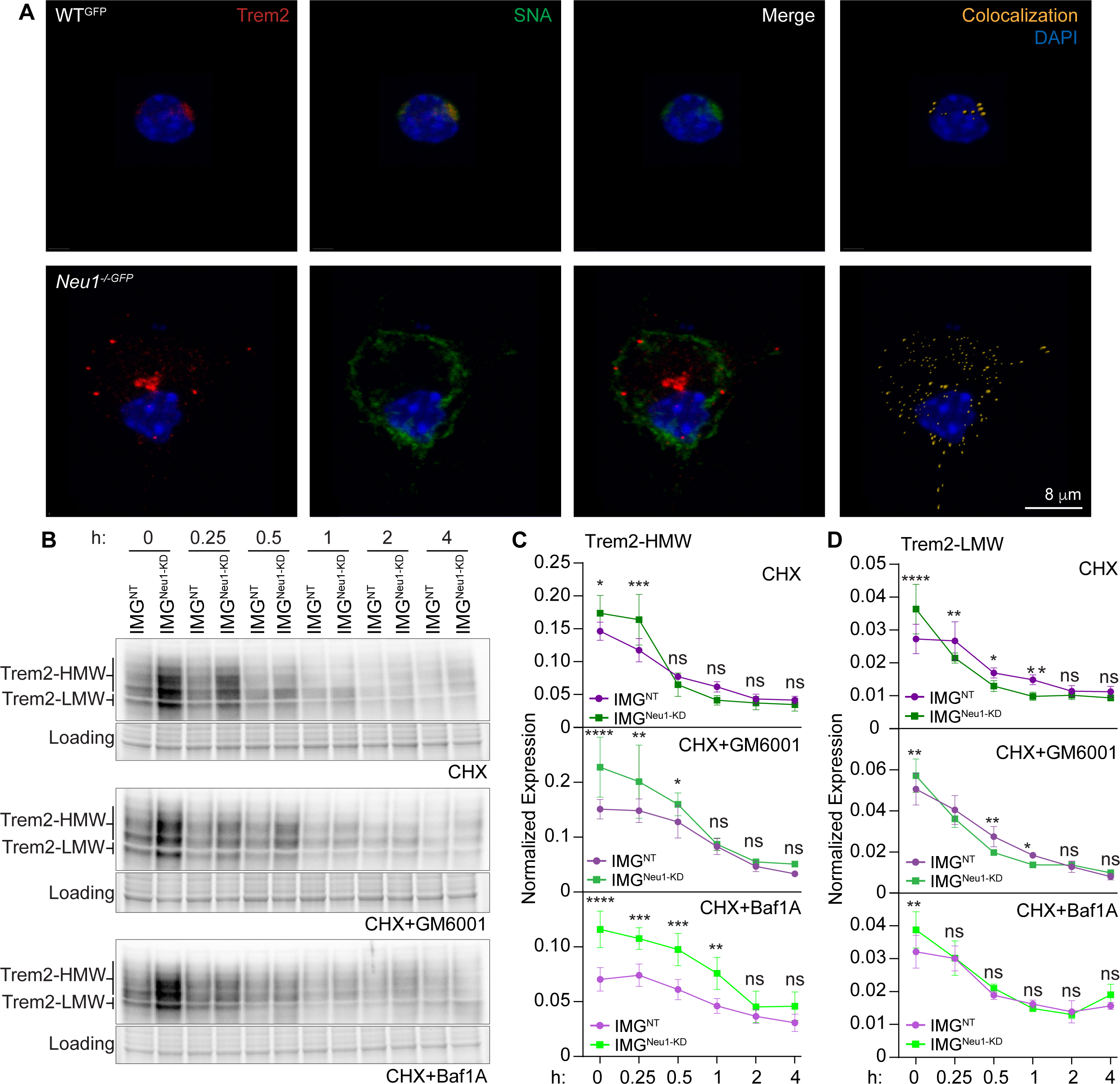
Expression of Sia–Trem2-FL in Neu1*^-/-^* microglia and after CHX treatment. (A) Representative images of IF staining showing sialylated Trem2 in 4-month-old WT^GFP^ and Neu1*^-/-^*^GFP^ microglia; Trem2 (red), SNA (green), DAPI (blue), and co-localized objects (yellow) are shown. (B) Representative immunoblots of Trem2-HMW and Trem2-LMW after treatment of IMG^NT^ and IMG^Neu1-KD^ cells with CHX only (top), CHX+GM6001 (middle), or CHX+Baf1A (bottom) for 0, 0.25, 0.5, 1, 2, and 4 h. (C-D) Normalized expression of (C) Trem2-HMW and (D) Trem2-LMW in IMG^NT^ and IMG^Neu1-KD^ cells after treatment with CHX only (top), CHX+GM6001 (middle), or CHX+Baf1A (bottom) for 0, 0.25, 0.5, 1, 2, and 4 h; n=5 for CHX-only groups, and n=4 for CHX+GM6001 and CHX+Baf1A groups. Data represented as the mean ± SEM; ns, not significant; *p≤0.05, **p≤0.01, ***p≤0.001, ****p≤0.0001. Related to Figure 5.

**Figure S4.**
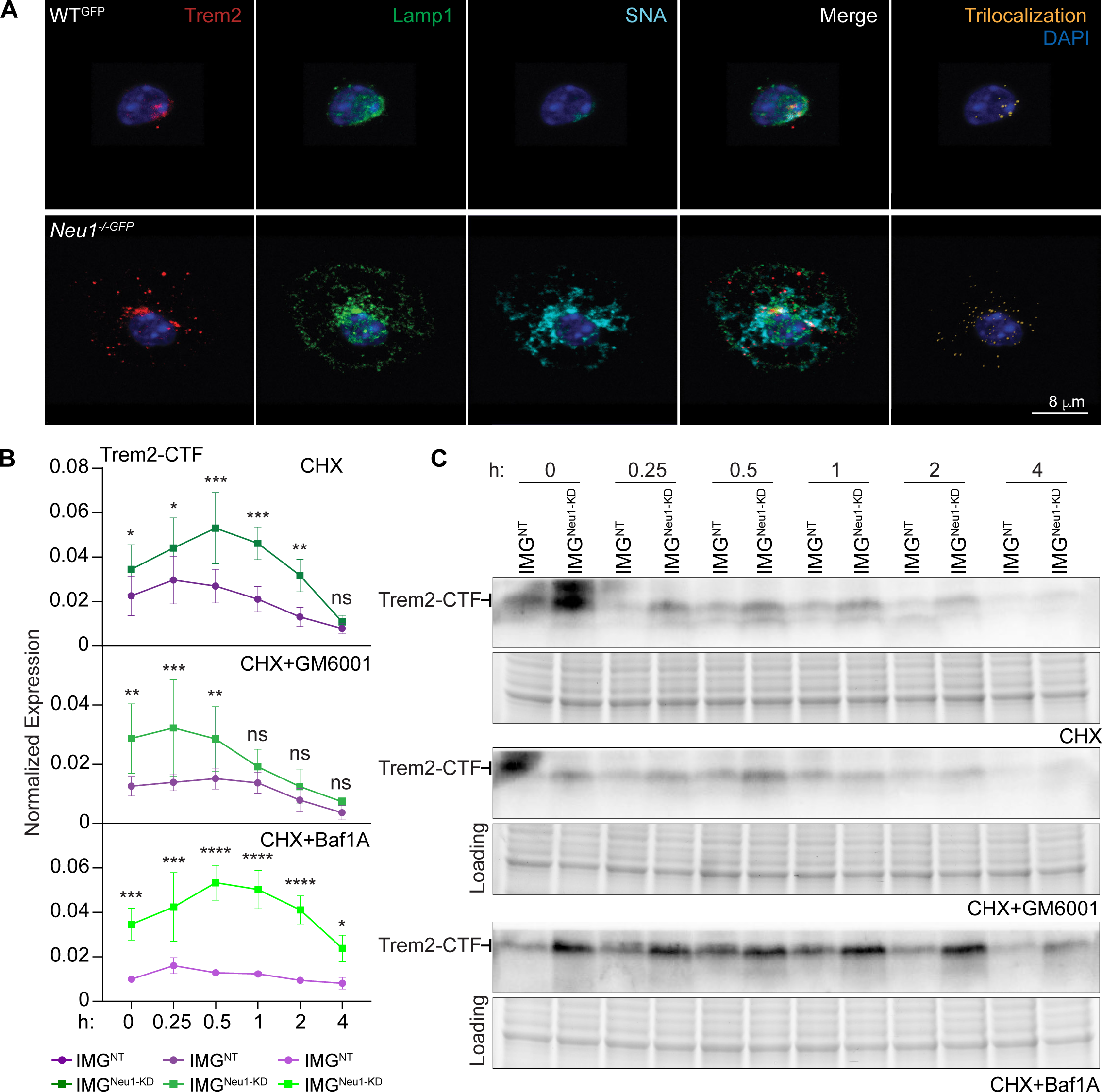
Lysosomal Sia–Trem2-FL in Neu1*^-/-^* microglia and Trem2-CTF expression after CHX treatment. (A) Representative images of IF staining showing lysosome-localized sialylated Trem2 in 4-month-old WT^GFP^ and Neu1*^-/-^*^GFP^ microglia. Trem2 (red), Lamp1 (green), SNA (cyan), DAPI (blue), and trilocalized objects (yellow) are shown. (B) Normalized expression of Trem2-CTF in IMG^NT^ and IMG^Neu1-KD^ cells after treatment with CHX only (top), CHX+GM6001 (middle), or CHX+Baf1A (bottom) for 0, 0.25, 0.5, 1, 2, and 4 h (n=5 for the CHX-only group, and n=4 for the CHX+GM6001 and CHX+Baf1A groups). Data represented as the mean ± SEM; ns, not significant; *p≤0.05, **p≤0.01, ***p≤0.001, ****p≤0.0001. (C) Representative immunoblots of Trem2-CTF in IMG^NT^ and IMG^Neu1-KD^ cells after treatment with CHX only (top), CHX+GM6001 (middle), or CHX+Baf1A (bottom). Related to Figure 5.

**Figure S5.**
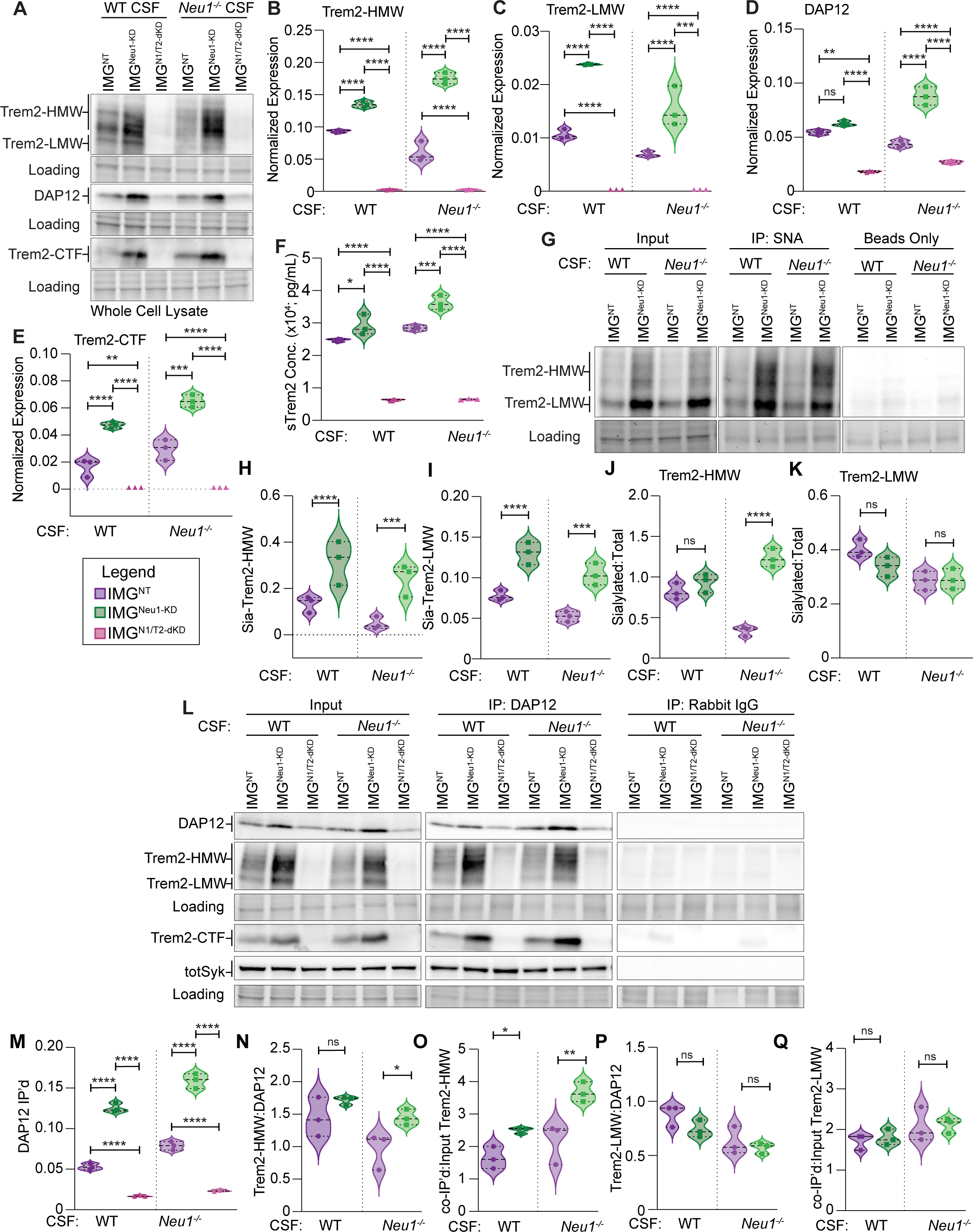
Sialylation of Trem2-HMW influences its association with DAP12 under homeostatic and inflammatory conditions. (A-Q) All samples were stimulated with 4-month-old WT or Neu1*^-/-^* CSF for 24 h. (A) Representative immunoblots of Trem2-HMW, Trem2-LMW, DAP12, and Trem2-CTF in stimulated IMG^NT^, IMG^Neu1-KD^, and IMG^N1/T1-dKD^ whole-cell lysates. (B-E) Quantification of (B) Trem2-HMW, (C) Trem2-LMW, (D) DAP12, and (E) Trem2-CTF in stimulated IMG^NT^, IMG^Neu1-KD^, and IMG^N1/T1-dKD^ whole-cell lysates. (F) Concentration of sTrem2 in the media of stimulated IMG^NT^, IMG^Neu1-KD^, and IMG^N1/T1-dKD^ cells. (G) Representative immunoblots of Trem2-HMW and Trem2-LMW in stimulated IMG^NT^ and IMG^Neu1-KD^ whole-cell lysates after immunoprecipitation (IP) of SNA; Input (left), SNA IP (middle), Beads-only control (right). (H-I) The ratio of total sialylated protein that is (H) Trem2-HMW and (I) Trem2-LMW in stimulated IMG^NT^ and IMG^Neu1-KD^ whole-cell lysates. (J-K) The ratios of sialylated (J) Trem2-HMW and (K) Trem2-LMW to total Trem2 in stimulated IMG^NT^ and IMG^Neu1-KD^ whole-cell lysates. (L) Representative immunoblots of DAP12, Trem2-HMW, Trem2-LMW, Trem2-CTF, and total Syk in stimulated IMG^NT^, IMG^Neu1-KD^, and IMG^N1/T2-dKD^ whole-cell lysates after immunoprecipitation of DAP12; Input (left), DAP12 IP (middle), Rabbit IgG control (right). (M) The normalized level of DAP12 that was immunoprecipitated from stimulated IMG^NT^, IMG^Neu1-KD^, and IMG^N1/T2-dKD^ whole-cell lysates. (N) The ratio of DAP12 that is in complex with Trem2-HMW in stimulated IMG^NT^ and IMG^Neu1-KD^ whole-cell lysates. (O) The ratio of Trem2-HMW that is in complex with DAP12 in stimulated IMG^NT^ and IMG^Neu1-KD^ whole-cell lysates. (P) The ratio of DAP12 that is in complex with Trem2-LMW in stimulated IMG^NT^ and IMG^Neu1-KD^ whole-cell lysates. (Q) The ratio of Trem2-LMW that is in complex with DAP12 in stimulated IMG^NT^ and IMG^Neu1-KD^ whole-cell lysates. (B-F, H-K, M-Q) The n=3 for all groups. Data represented as the medians and quartiles; ns, not significant; *p≤0.05, **p≤0.01, ***p≤0.001, ****p≤0.0001. Related to Figure 6.

**Figure S6.**
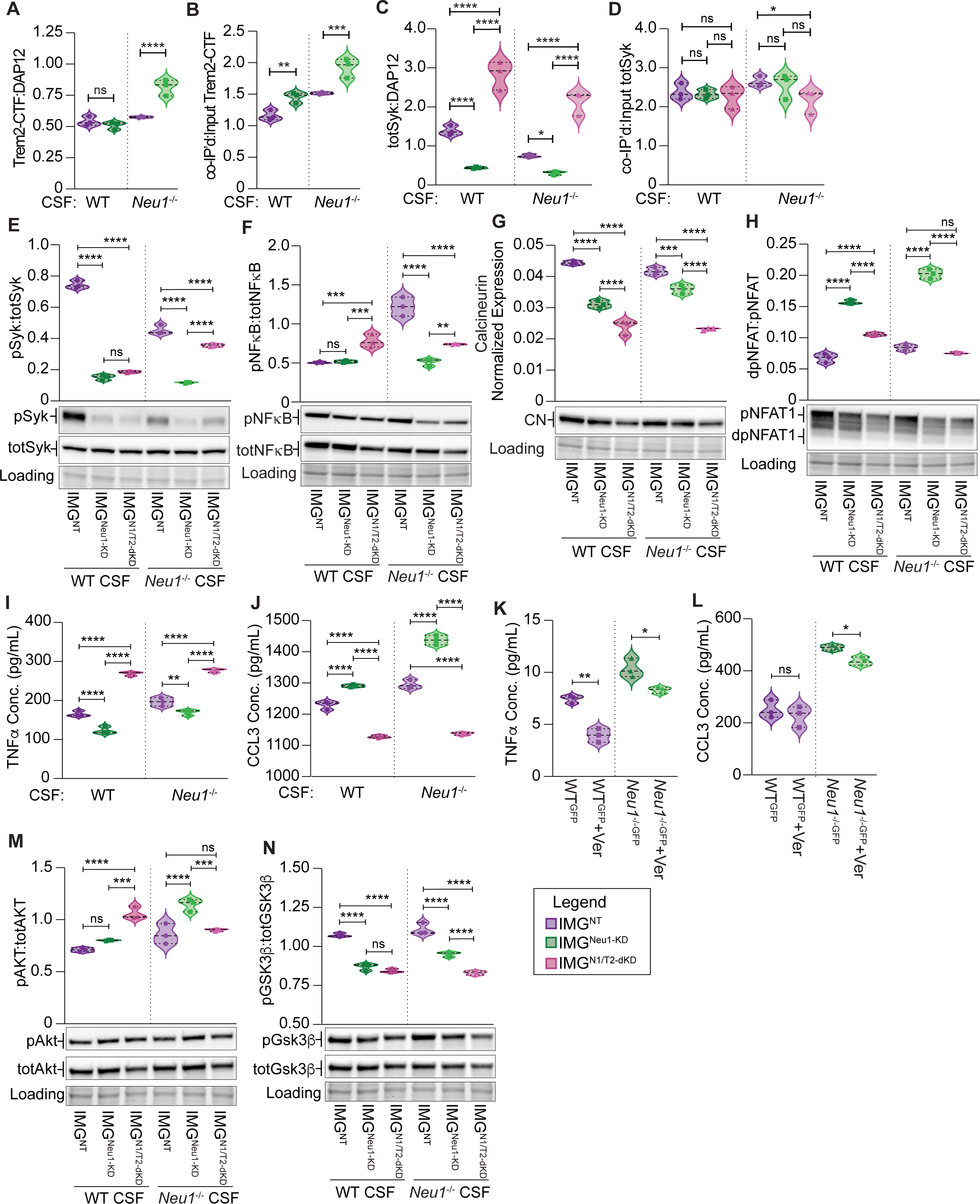
Downregulation of Neu1 enzymatic activity alters microglial signaling downstream of Trem2 activation. (A-N) All samples were stimulated with 4-month-old WT or Neu1*^-/-^* CSF for 24 h. (A-D) All results were obtained after immunoprecipitation (IP) of DAP12. (A) The ratio of DAP12 that is in complex with Trem2-CTF in stimulated IMG^NT^ and IMG^Neu1-KD^ whole-cell lysates. (B) The ratio of Trem2-CTF that is in complex with DAP12 in stimulated IMG^NT^ and IMG^Neu1-KD^ whole-cell lysates. (C) The ratio of DAP12 that is in complex with total Syk in stimulated IMG^NT^, IMG^Neu1-KD^, and IMG^N1/T2-dKD^ whole-cell lysates. (D) The ratio of total Syk that is in complex with DAP12 in stimulated IMG^NT^, IMG^Neu1-KD^, and IMG^N1/T2-dKD^ whole-cell lysates. (E-H) Representative immunoblots and the activation of signaling molecules represented as the (E) ratio of pSyk:totSyk, (F) ratio of pNF*k*B:totNF*k*B, (G) normalized expression of calcineurin (CN), (H) ratio of dpNFAT1:pNFAT1 in stimulated IMG^NT^, IMG^Neu1-KD^, and IMG^N1/T2-dKD^ cells. (I-J) Concentrations of (I) TNF*a* and (J) CCL3 in media from stimulated IMG^NT^, IMG^Neu1-KD^, and IMG^N1/T2-dKD^ cells. (K-L) Concentrations of (K) TNF*a* and (L) CCL3 in media from primary 4-month-old WT^GFP^ and Neu1*^-/-^*^GFP^ microglia maintained in culture with or without 50 *μ*M verapamil (Ver) for 4 h. (M-N) Representative immunoblots and the activation of signaling molecules represented as the (M) ratio of pAkt:totAkt, and (N) ratio of pGsk3*b*:totGsk3*b* in stimulated IMG^NT^, IMG^Neu1-KD^, and IMG^N1/T2-dKD^ cells. The n=3 for all groups, and data represented as median and quartiles; ns, not significant; *p≤0.05, **p≤0.01, ***p≤0.001, ****p≤0.0001. Related to Figure 6.

**Figure S7.**
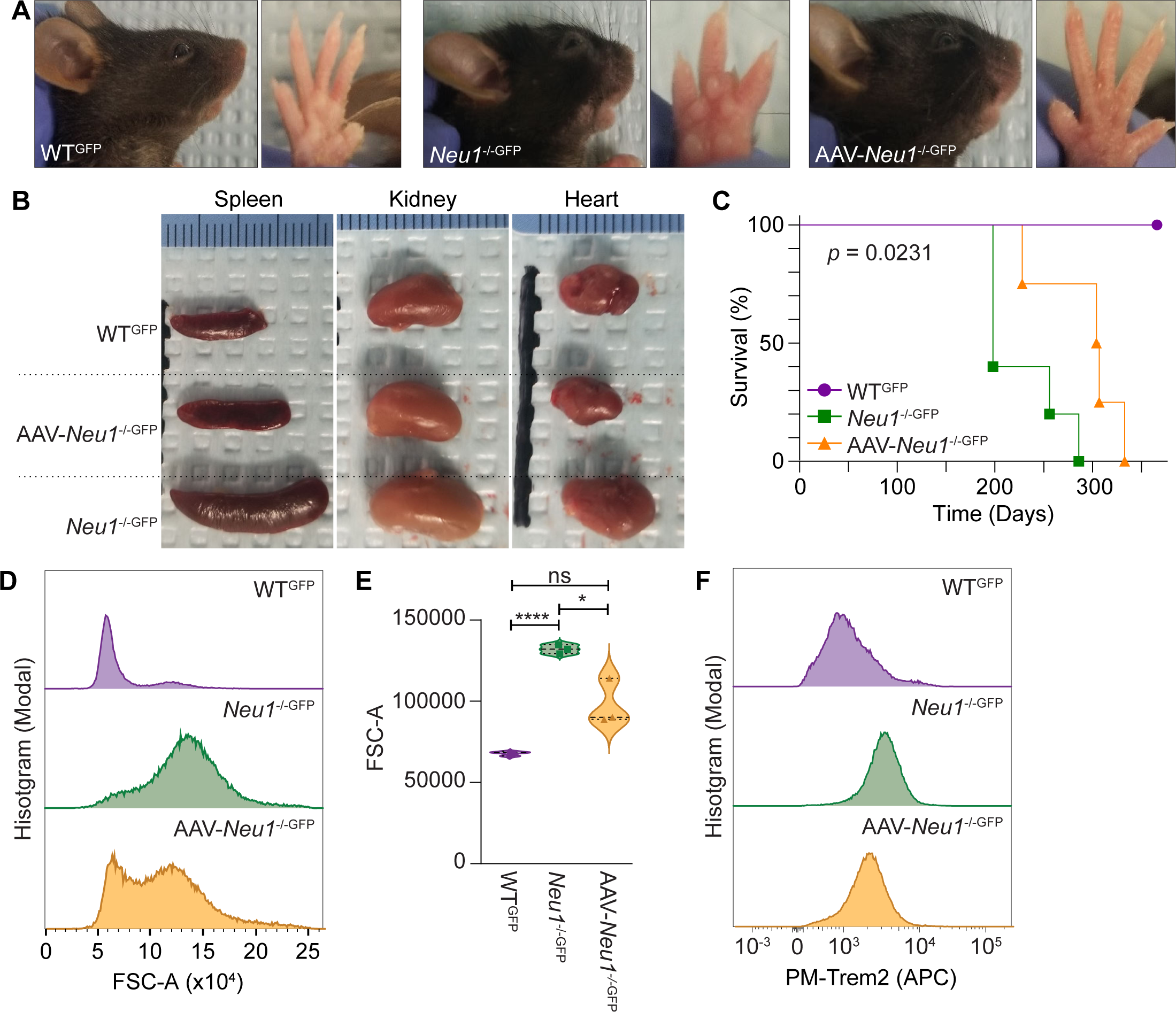
Reinstatement of Neu1 enzymatic activity corrects pathologic features of sialidosis and improves survival. (A) Representative photographs of facial morphology and edema in the paws of 4-month-old WT^GFP^, Neu1*^-/-^*^GFP^, and AAV-Neu1*^-/-^*^GFP^ mice. (B) Representative photographs showing the sizes of the spleen, kidney, and heart of 4-month-old WT^GFP^, Neu1*^-/-^*^GFP^, and AAV-Neu1*^-/-^*^GFP^ mice. (C) Survival of WT^GFP^, Neu1*^-/-^*^GFP^, and AAV-Neu1*^-/-^*^GFP^ mice during the 12-month treatment period; n=5 for WT^GFP^ and Neu1*^-/-^*^GFP^ mice, and n=4 for AAV-Neu1*^-/-^*^GFP^ mice; p=0.0231. (D) Representative plots showing the cell size (FSC-A) of 4-month-old WT^GFP^, Neu1*^-/-^*^GFP^, and AAV-Neu1*^-/-^*^GFP^ microglia. (E) Quantification of cell size (FSC-A) of 4-month-old WT^GFP^, Neu1*^-/-^*^GFP^, and AAV-Neu1*^-/-^*^GFP^ microglia, as measured by FACS; n=3 for all samples. Data represented as the medians and quartiles; ns, not significant; *p≤0.05, ****p≤0.0001. (F) Representative plots showing PM-Trem2-FL expression in 4-month-old WT^GFP^, Neu1*^-/-^*^GFP^ and AAV-Neu1*^-/-^*^GFP^ microglia. Related to Figure 7.

**Table S1.**
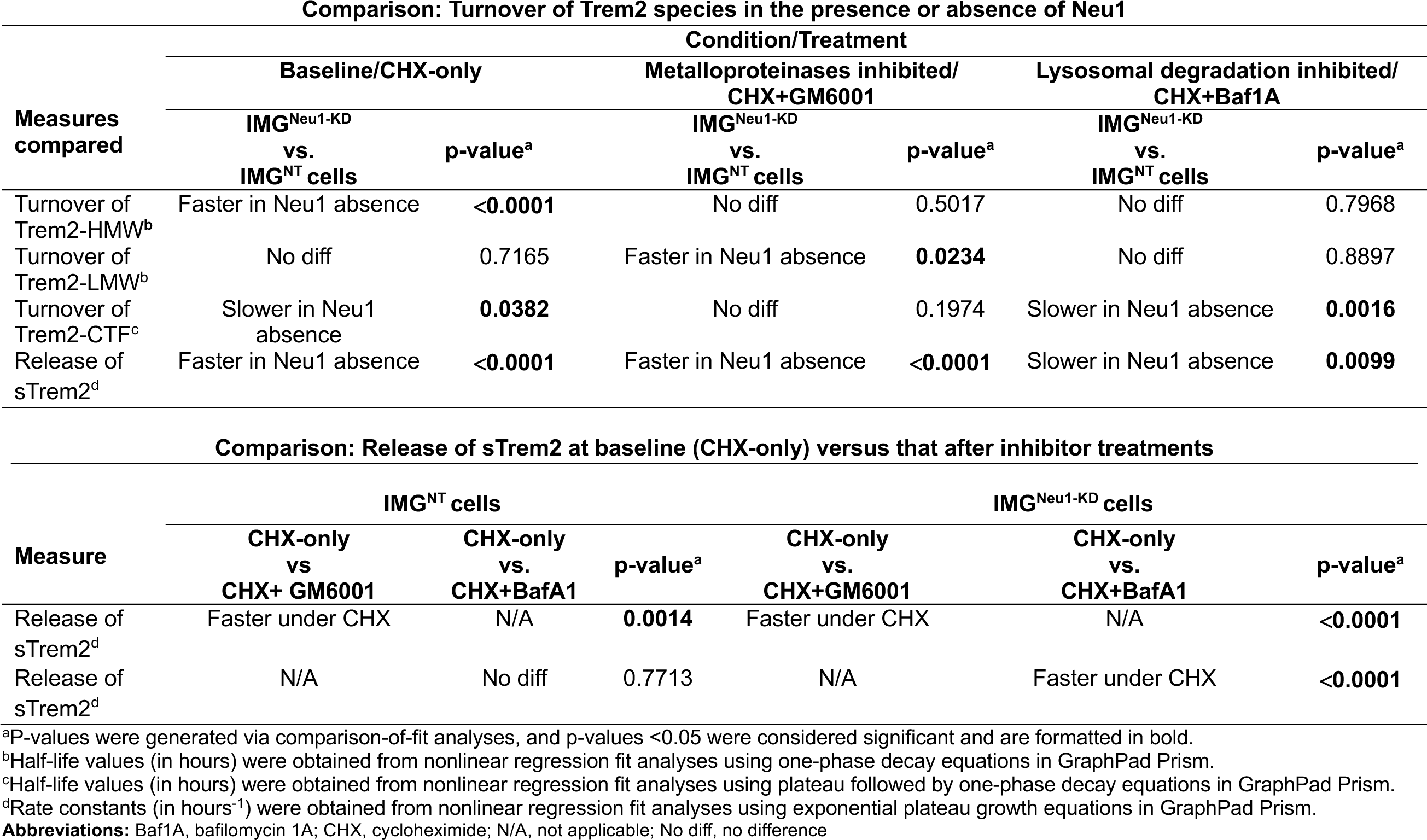
Comparisons of Trem2 species turnover in IMG^NT^ cells vs. IMG^Neu1-KD^ cells. Related to Figures 5, S3, and S4.

| **Comparison: Turnover of Trem2 species in the presence or absence of Neu1** |  |  |  |  |  |  |
| --- | --- | --- | --- | --- | --- | --- |
| **Measures compared** | **Condition/Treatment** |  |  |  |  |  |
|  | **Baseline/CHX-only** |  | **Metalloproteinases inhibited/**<br>**CHX+GM6001** |  | **Lysosomal degradation inhibited/**<br>**CHX+Baf1A** |  |
|  | **IMG<sup>Neu1-KD</sup>**<br>**vs.**<br>**IMG<sup>NT</sup> cells** | **p-value<sup>a</sup>** | **IMG<sup>Neu1-KD</sup>**<br>**vs.**<br>**IMG<sup>NT</sup> cells** | **p-value<sup>a</sup>** | **IMG<sup>Neu1-KD</sup>**<br>**vs.**<br>**IMG<sup>NT</sup> cells** | **p-value<sup>a</sup>** |
| Turnover of Trem2-HMW<sup>b</sup> | Faster in Neu1 absence | **<0.0001** | No diff | 0.5017 | No diff | 0.7968 |
| Turnover of Trem2-LMW<sup>b</sup> | No diff | 0.7165 | Faster in Neu1 absence | **0.0234** | No diff | 0.8897 |
| Turnover of Trem2-CTF<sup>c</sup> | Slower in Neu1 absence | **0.0382** | No diff | 0.1974 | Slower in Neu1 absence | **0.0016** |
| Release of sTrem2<sup>d</sup> | Faster in Neu1 absence | **<0.0001** | Faster in Neu1 absence | **<0.0001** | Slower in Neu1 absence | **0.0099** |
| **Comparison: Release of sTrem2 at baseline (CHX-only) versus that after inhibitor treatments** |  |  |  |  |  |  |
| **Measure** | **IMG<sup>NT</sup> cells** |  |  | **IMG<sup>Neu1-KD</sup> cells** |  |  |
|  | **CHX-only**<br>**vs.**<br>**CHX+ GM6001** | **CHX-only**<br>**vs.**<br>**CHX+BafA1** | **p-value<sup>a</sup>** | **CHX-only**<br>**vs.**<br>**CHX+GM6001** | **CHX-only**<br>**vs.**<br>**CHX+BafA1** | **p-value<sup>a</sup>** |
| Release of sTrem2<sup>d</sup> | Faster under CHX | N/A | **0.0014** | Faster under CHX | N/A | **<0.0001** |
| Release of sTrem2<sup>d</sup> | N/A | No diff | 0.7713 | N/A | Faster under CHX | **<0.0001** |

<sup>a</sup>P-values were generated via comparison-of-fit analyses, and p-values <0.05 were considered significant and are formatted in bold.<sup>b</sup>Half-life values (in hours) were obtained from nonlinear regression fit analyses using one-phase decay equations in GraphPad Prism.<sup>c</sup>Half-life values (in hours) were obtained from nonlinear regression fit analyses using plateau followed by one-phase decay equations in GraphPad Prism.<sup>d</sup>Rate constants (in hours<sup>-1</sup>) were obtained from nonlinear regression fit analyses using exponential plateau growth equations in GraphPad Prism.**Abbreviations:** Baf1A, bafilomycin 1A; CHX, cycloheximide; N/A, not applicable; No diff, no difference

**Table S2.** Pro-inflammation affects the sialylation of Term2 species in IMG^NT^ cells and IMG^Neu1-KD^ cells. Related to Figure 6.

| Trem2<br>Species<br>Measured<sup>c</sup> | IMG<sup>NT</sup> cells<sup>a</sup> |  | IMG<sup>Neu1-KD</sup> cells<sup>a</sup> |  |
| --- | --- | --- | --- | --- |
|  | Homeostasis<br>vs.<br>Pro-inflammation | p-value<sup>b</sup> | Homeostasis<br>vs.<br>Pro-inflammation | p-value<sup>b</sup> |
| Sia–Trem2-HMW | Decreased under pro-inflammation | **0.0007** | Increased under pro-inflammation | **0.0042** |
| Sia–Trem2-LMW | Decreased under pro-inflammation | **0.0188** | No difference | 0.1899 |

<sup>a</sup> Cells were treated with either WT (homeostatic) or *Neu1*<sup>-/-</sup> (pro-inflammatory) CSF.
<sup>b</sup> P-values were generated using 2-way ANOVA analyses; bold formatting indicates significant differences between treatments.
**Abbreviations:** CSF, cerebrospinal fluid; HMW, higher molecular weight; LMW, low molecular weight; Sia, sialylated

**Table S3.** Neu1 affects the interactions between some Trem2 species and DAP12 during homeostasis and pro-inflammatory conditions. Related to Figure 6.

| Interactions | IMG<sup>NT</sup> Cells |  |  | IMG<sup>Neu1-KD</sup> Cells |  |  |
| --- | --- | --- | --- | --- | --- | --- |
|  | WT CSF<sup>a</sup> | Neu1<sup>-/-</sup> CSF<sup>b</sup> | p-value<sup>c</sup> | WT CSF<sup>a</sup> | Neu1<sup>-/-</sup> CSF<sup>b</sup> | p-value<sup>c</sup> |
| DAP12 complexed with Trem2-HMW<sup>d</sup> | 1.445 ± 0.301 | 0.962 ± 0.278 | **0.0392** | 1.714 ± 0.063 | 1.447 ± 0.122 | 0.1699 |
| DAP12 complexed with Trem2-LMW<sup>d</sup> | 0.879 ± 0.101 | 0.624 ± 0.128 | 0.0547 | 0.740 ± 0.080 | 0.572 ± 0.051 | 0.1512 |
| DAP12 complexed with Trem2-CTF<sup>d</sup> | 0.541 ± 0.038 | 0.575 ± 0.006 | 0.4338 | 0.510 ± 0.030 | 0.816 ± 0.065 | **0.0015** |
| Trem2-HMW complexed with DAP12<sup>e</sup> | 1.642 ± 0.342 | 2.166 ± 0.626 | 0.1481 | 2.501 ± 0.092 | 3.664 ± 0.308 | **0.0165** |
| Trem2-LMW complexed with DAP12<sup>e</sup> | 1.715 ± 0.197 | 2.076 ± 0.427 | 0.2337 | 1.796 ± 0.194 | 2.118 ± 0.188 | 0.2782 |
| Trem2-CTF complexed with DAP12<sup>e</sup> | 1.161 ± 0.081 | 1.516 ± 0.017 | **0.0219** | 1.453 ± 0.085 | 1.926 ± 0.152 | **0.0083** |

<sup>a</sup>WT CSF treatment mimics the homeostatic microenvironment. Values represent mean ± SD.
<sup>b</sup>Neu1<sup>-/-</sup> CSF treatment induces a pro-inflammatory microenvironment. Values represent mean ± SD.
<sup>c</sup>Two-way ANOVA analyses were done to calculate p-values. P < 0.05 (in bold) indicates a significant difference in interactions between the two treatments.
<sup>d</sup>This interaction notes the ratio of the amount of the Trem2 species co-immunoprecipitated with DAP12 antibodies to the total amount of DAP12 immunoprecipitated.
<sup>e</sup>This interaction notes the ratio of the amount of the Trem2 species co-immunoprecipitated with DAP12 antibodies to the total input of that Trem2 species.

**Table S4.** Concentrations of Trem2 species, DAP12, and Trem2-signaling molecules are altered in different microenvironments. Related to Figure 6.

| Factor measured | Cell Lines |  |  |  |  |  |
| --- | --- | --- | --- | --- | --- | --- |
|  | IMG<sup>NT</sup> |  | IMG<sup>Neu1-KD</sup> |  | IMG<sup>N1/T2-dKD</sup> |  |
|  | Level change under pro-inflammation vs. homeostasis<sup>a</sup> | p-value<sup>b</sup> | Level change under pro-inflammation vs. homeostasis<sup>a</sup> | p-value<sup>b</sup> | Level change under pro-inflammation vs. homeostasis<sup>a</sup> | p-value<sup>b</sup> |
| Trem2-HMW | Decrease | **0.0007** | Increase | **0.0002** | Not present | >0.9999 |
| Trem2-LMW | Decrease | **0.0320** | Decrease | **0.0008** | Not present | >0.9999 |
| DAP12 | Decrease | **0.0081** | Increase | **0.0001** | Increase | **0.0211** |
| Trem2-CTF | Increase | **0.0166** | Increase | **0.0040** | No present | >0.9999 |
| sTrem2 | Increase | **0.0002** | Increase | **<0.0001** | No difference | 0.6842 |
| Activated Syk | Decrease | **<0.0001** | No difference | 0.0727 | Increase | **<0.0001** |
| Activated NFkB | Increase | **<0.0001** | No difference | 0.9135 | No difference | 0.4810 |
| Calcineurin | Decrease | **0.0336** | Increase | **0.0024** | No difference | 0.4806 |
| Activated NFAT1 | Increase | **0.0172** | Increase | **0.0001** | Decrease | **0.0010** |
| Activated Akt | Increase | **0.0113** | Increase | **0.0002** | Decrease | **0.0086** |
| Inhibited GSK3β | No difference | 0.0956 | Increase | **0.0048** | No difference | 0.5130 |

<sup>a</sup>Pro-inflammation refers to cells treated with *Neu1*<sup>-/-</sup> CSF, and homeostasis refers to cells treated with wild-type CSF.
<sup>b</sup>P-values were calculated by 2-way ANOVA analysis, and p <0.05 was considered significant (formatted in bold).

**Table S5.** Neu1 downregulation in neurological conditions, diseases, and disorders^a^, related to Discussion.

| Category | Disease or population | Sample Type | NEU1 log-fold change<sup>b</sup> | P-value | Dataset ID | Testing platform | Original publication |
| --- | --- | --- | --- | --- | --- | --- | --- |
| **Neurodegeneration** | HIV with encephalitis | Brain, frontal cortex | −0.828 | 0.00174 | GSE35864 | Affymetrix HG-U133_Plus_2 | Gelman BB, et. al., *PLoS One* 2012. PMID: 23049970 |
|  | Multiple sclerosis | PBMCs | −0.478 | 0.00012 | GSE21942 | Affymetrix HG-U133_Plus_2 | Kemppinen AK, et. al., *BMJ Open* 2011. PMID: 22021740 |
|  | FTLD-U with progranulin mutations | Brain, hippocampus | −1.251 | 0.00156 | GSE13126 | Affymetrix HG-133A_2 | Chen-Plotkin AS, et. al., *Hum Mol Genet* 2008. PMID: 18223198 |
|  | Alzheimer disease | Brain, hippocampus | −0.295 | 0.00105 | GSE36980 | Affymetrix HuGene-1_0-st | Hokama M, et. al., *Cereb Cortex* 2014. PMID: 23595620 |
|  |  | Brain, hippocampus | −0.449 | 0.08692 | GSE173955 | Illumina HiSeq 1500 | Mizuno Y, et. al., *Oxid Med Cell Longev* 2021. PMID: 34970419 |
|  |  | Brain, hippocampus-CA1 region | −1.173 | 0.03590 | GSE1297 | Affymetrix HG-133A_2 | Blalock EM, et. al., *Proc Natl Acad Sci U S A* 2004. PMID: 14769913 |
| **Neuropsychiatric** | Schizophrenia | Brain, prefrontal cortex-BA10 | −0.313 | 0.04500 | GSE12654 | Affymetrix HG_U95Av2 | Iwamoto K, et. al., *Mol Psychiatry* 2004. PMID: 14743183 |
|  | Bipolar disorder | Brain, dorsolateral prefrontal cortex | −0.164 | 0.04250 | GSE5388 | Affymetrix HG-133A_2 | Ryan MM, et. Al., *Mol Psychiatry* 2006. PMID: 16894394 |
| **Neurodevelopmental** | Autism | Peripheral blood lymphocytes | −0.121 | 0.03970 | GSE25507 | Affymetrix HG-U133_Plus_2 | Alter MD, et. al., *PloS One* 2011. PMID: 21379579 |
| Substance abuse | Cocaine | Brain, midbrain | −0.121 | 0.02420 | GSE4839 | Illumina HumanHT-12 V3.0 | Bannon MJ, et. al., *Neuropsychopharmacology* 2014. PMID: 24642598 |
|  | Alcohol | Brain, hippocampus | −0.201 | 0.04300 | GSE44456 | Affymetrix HuGene-1_0-st | McClintick JN, et. al., *Alcohol* 2013. PMID: 23981442 |
| Aging | <70 y vs. >70 y | Brain, frontal cortex | −0.221 | 0.01060 | GSE53890 | Affymetrix HG-U133_Plus_2 | Lu T, et. al., *Nature* 2014. PMID: 24670762 |
|  |  | Brain, frontal cortex | −0.225 | 0.11000 | GSE1572 | Affymetrix HG_U95Av2 | Lu T, et. al., *Nature* 2004. PMID: 15190254 |

<sup>a</sup>Data were obtained from the publicly available GEO DataSets from NCBI. Information for each data set includes the category, patient/tested population or disease, sample type, log-fold change of *NEU1* expression, *p*-value, dataset ID, testing platform, and original publication citation.
<sup>b</sup>*NEU1* expression was calculated as the log-fold change of *NEU1* RNA in tested groups vs. control groups by using GEO2R.
**Abbreviations:** FTLD-U, frontotemporal lobar degeneration with ubiquitin<sup>+</sup> inclusions; GSE, Genomic Spatial Event database; PBMCs, peripheral blood mononuclear cells

**Table S6.**
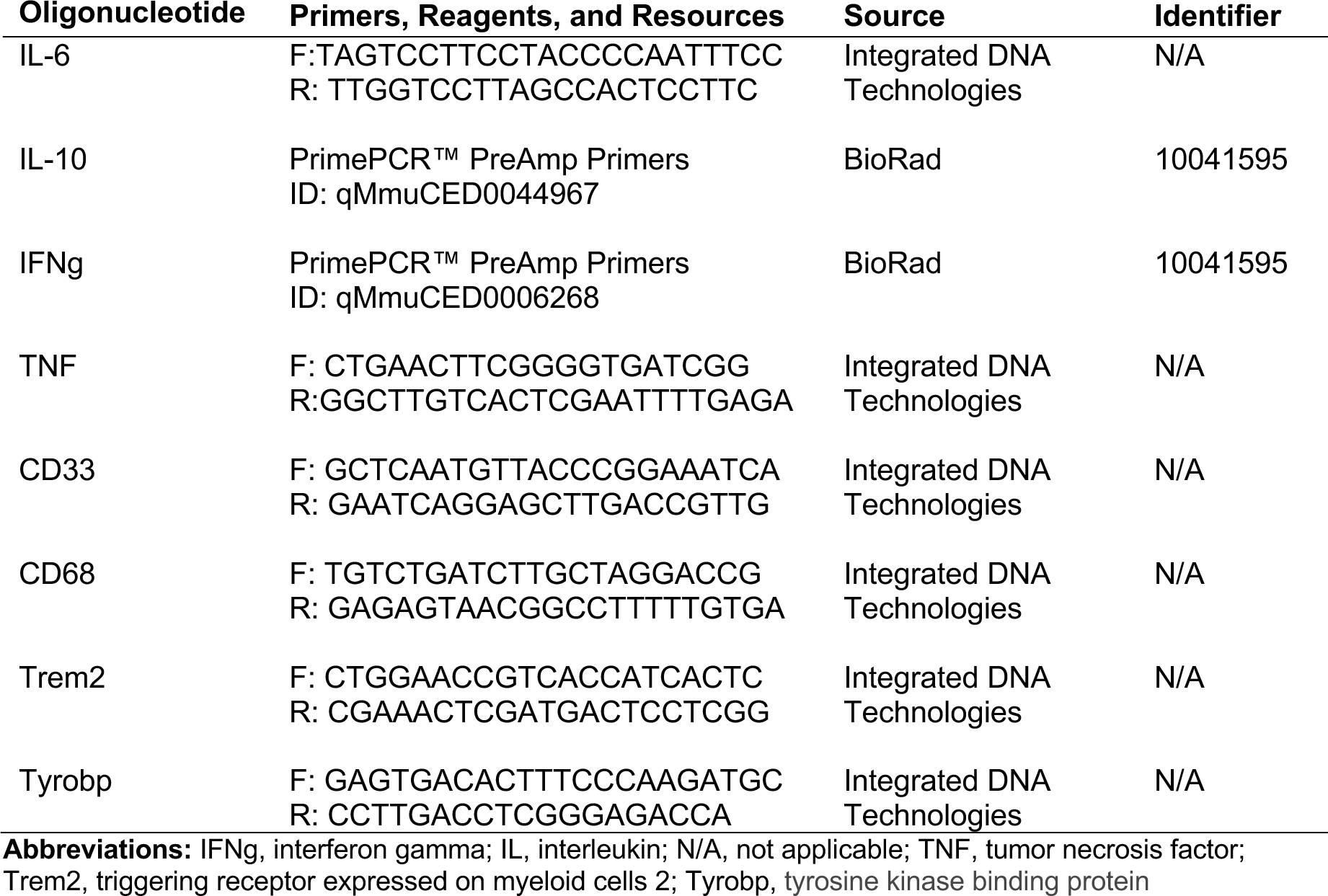
Resource table continued: list of oligonucleotides, related to STAR Methods.

## Notes

### Competing Interest Statement

The authors have declared no competing interest.

